# Laboratory application of the Aquascope approach of automated imaging and classification for long-term plankton monitoring

**DOI:** 10.1101/2024.02.23.581739

**Authors:** Stefanie Merkli, Marta Reyes, Francesco Pomati

## Abstract

Anthropogenic changes such as climate change and pollution have strong effects on plankton and its ecosystem services. In freshwater, phytoplankton forms the basis of the food web and plays a key role in water quality. It is therefore important to monitor the plankton community. In Switzerland, lake plankton has been monitored for decades by cantonal authorities. But in the future, there will be a limitation in the availability of trained taxonomists that are able to count plankton samples under the microscope as well as time and budget constraints. New technologies will be needed to fulfil the federal requirements on the monitoring of lakes in the future. Here we focus on two such instruments, both based on plankton automated imaging: the FlowCam and the Aquascope. The first is a commercially available instrument, the second is custom made. They are the only automated imaging instruments that can track freshwater plankton across the desired size range (phytoplankton and zooplankton). Their design makes them the state of the art technologies to make quantitative observations of plankton. While the FlowCam is a laboratory instrument, the Aquascope is originally designed for field deployment. In this report, we focus the empirical work on the laboratory application of the Aquascope approach of automated imaging and classification for long term plankton monitoring. We find that there are major advantages of the Aquascope approach but the application as a laboratory instrument still needs further development. The major advantages of the Aquascope approach compared to traditional, human-supervised, microscopy methods are the automation and the speed of the data processing, the availability of individual level traits and the management of the data generated. One of the major drawbacks is the lack of taxonomic resolution (mostly at the genus level). Nevertheless, we show that with a larger benchmarking project, it might be possible to continue the long term time series on a coarse taxonomic level as usually reported by cantonal authorities to the general public. We discuss advantages and limitations of Aquascope relative to traditional microscopy and FlowCam, and propose ways forward for future development and application of this approach.

## Introduction

### Status quo of Plankton Monitoring Switzerland

Anthropogenic changes such as climate change and pollution have strong effects on plankton and its ecosystem services (e.g. drinking water, oxygen production, fish yield, etc.) (Bundesamt für Umwelt BAFU 2022). Phytoplankton cover an enormous phylogenetic and functional diversity, and drive essential global scale processes and elemental cycles, such as those of carbon and nutrients, which are and have been shaping Earth’s atmosphere chemistry for billions of years (Falkowski et al. 2004). In freshwater, phytoplankton forms the basis of the food web and plays a key role in water quality (Lombard et al. 2019). It is therefore important to monitor the plankton community.

In Switzerland, lake plankton has been monitored regularly for decades. “The Water Protection Act (Art. 50 GSchG) obliges the Confederation and the cantons to examine the effects of water protection measures and to inform the public about the condition of water bodies. The federal government is responsible for the surveys (Art. 57 GSchG) and the information (Art. 49 GSchV), which are of nationwide interest. The cantons carry out all other surveys required for the enforcement of the Water Protection Act (Art. 58 GSchG) and are responsible for providing information on the state of water bodies and water protection in their cantonal territory (Art. 49 GSchV). They are also obliged to report their findings to the federal authorities. If the water quality does not meet the requirements, the nature, extent and causes of the pollution must be determined (Art. 47 GSchV) and the effectiveness of measures must be assessed.” (Niederhauser 2023).

Traditionally, plankton samples are counted manually under the microscope. This technique requires a lot of experience and time from trained taxonomists. The quality of the analysis is therefore strongly affected by the taxonomists and mistakes are highly variable between people and not traceable (Merz et al. 2021; Álvarez et al. 2013; Rivas-Villar et al. 2021, 2020). Counts are then related to standard biovolumes of a given taxon. Biovolume standards do not account for individual-level size differences, as well as changes in size across the season, across decades or across different lakes. For zooplankton, standard sizes are not able to grasp the size distribution between juvenile and adult stages. This size resolution biases the estimation of the overall phytoplankton and zooplankton biomass in a given lake.

Further, lakes vary largely in their size, depth and eutrophication state which affects the sampling method. Other factors such as financial and time constraints and historic sampling effort led to differences between monitoring campaigns in different cantons. As there will be a limitation in the availability of trained taxonomists that are able to count plankton samples as well as future time and budget constraints, new technologies will be needed to fulfil the federal requirements on the monitoring of lakes in the near future. The needs and expectations for future plankton monitoring vary between cantons as much as their current methods. Here, we therefore discuss different methods for automated plankton monitoring that could be used to fill this future gap.

### State of the art automated plankton enumeration

In 2019, Lombard et al. reviewed the state of the art technologies to make quantitative observations of plankton. Most of the technologies are designed for use in the ocean, a much more challenging environment compared to freshwaters in terms of water physics, chemistry, meteorological conditions for deployment and diversity in terms of plankton taxa. Pigment or acoustic methods can only distinguish functional groups with little compositional information and genetic methods are still largely constrained by a lack of databases and the possibility of quantification of absolute abundance rather than relative abundances. Instruments that can distinguish and allow counting of taxa are limited to imaging techniques.

Lombard et al.’s Fig. 1 gives a good overview of the currently commercially available imaging instruments. For freshwater plankton, we are generally interested in particles from 1 μm to 1 cm (Merz et al. 2021).

**Fig. 1:**
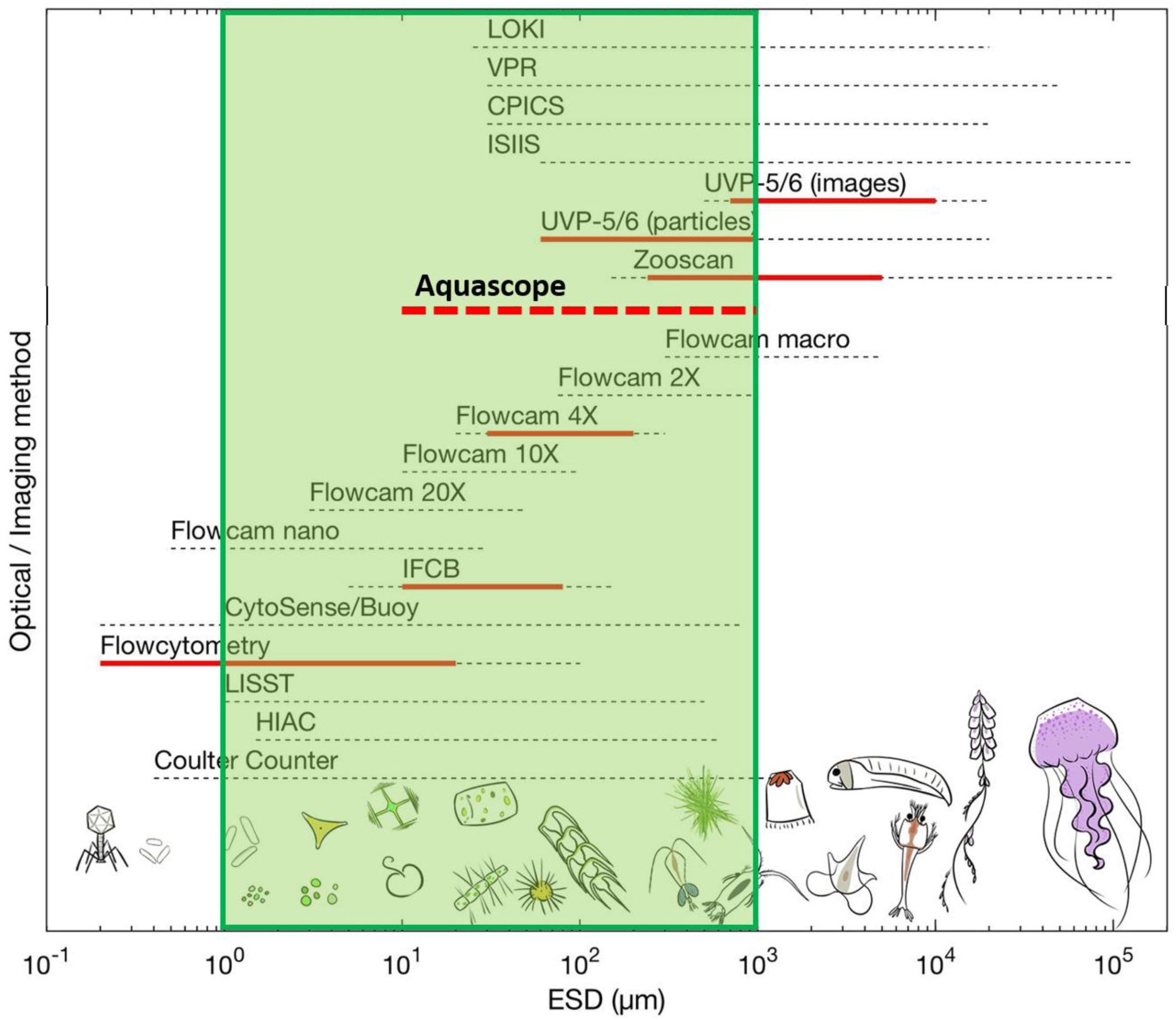
Adapted from Lombard et al. (2019): “Comparison of the total size range of plankton (in equivalent spherical diameter; ESD) that available optical and imaging methods can sample. Dashed lines represent the total operational size range from commercial information while the red line represents the practical size range which is efficient to obtain quantitative information […].” The green rectangle represents the size range of freshwater plankton. The dashed red line represents the detectable size range of the Aquascope.

An additional imaging system designed to image plankton and has not been covered by the 2019 review is the Scripps Plankton Camera (SPC) system. As described in Merz et al. (2021), Francesco Pomati and his team, in collaboration with the group of Jules Jaffe (Scripps Institution of Oceanography, UCSD, CA), adapted the SPC to a dual-magnification dark field underwater microscope (DSPC, in this report referred to as Aquascope) which is able to track particles from ∼ 10 μm to 1 cm. In the figure from Lombard et al. the Aquascope would lay in the range of the FlowCam 20X to the FlowCam 2X (**Fig. 1**). **With the FlowCam, the Aquascope is the only automated imaging instrument that can track freshwater plankton in the desired size range with only one machine.**

In this report we introduce three imaging techniques for plankton monitoring.

#### Microscopy

Rivas-Villar et al. suggested a method for automated image analysis similar to traditional microscopy counts (Rivas-Villar et al. 2021, 2020). They argue that, while classic monitoring is done with a microscope, images can be acquired by a lab-technician and then analysed using ML algorithms. They developed an algorithm that partitions plankton cells (and colonies) in microscopy images and then classifies those into taxonomic units. For this, a technician needs to manually image glutaraldehyde fixed samples under the microscope but the enumeration can be automated and the identification is traceable. They subject the sample to vacuum before sedimentation in order to explode vacuoles, which ensures the floating cyanobacteria to sink to the bottom of the (Utermöhl) counting chamber. According to Rivas-Villar et al. this technique is simple but it requires sampling, sample preparation, and there is still a need for further development. Glutaraldehyde samples require care and assessment of risks related to its toxicity. We assume that for this method, one would first have to somehow determine the density of the sample to ensure that cells do not overlap in the images taken for automated image detection. In addition, a ML classification for those images would have to be built from scratch.

#### FlowCam

An already established instrument (and method) is the FlowCam. The company page notes that “FlowCam was originally developed in the mid-1990s at Bigelow Laboratory for Ocean Sciences for studying plankton in ocean water. Commercialized by Fluid Imaging Technologies, FlowCam is designed to combine the benefits of digital imaging, flow cytometry, and microscopy into a single instrument. In April of 2020, Fluid Imaging Technologies was acquired by Yokogawa Electric Corporation, a leading provider of Industrial Automation and Test and Measurement solutions. The addition of FlowCam to the Life Innovation Business unit of Yokogawa supports the strategic expansion into the growing life sciences market.” (Yokogawa Fluid Imaging Technologies, Inc n.d.) According to the Yokogawa Electric Corporation, the target markets for the FlowCam are water & wastewater, food & beverage and pharmaceutical & medicine (Yokogawa Electric Corporation n.d.). They have also developed a FlowCam Cyano which is supposed to be able to detect phycocyanin and chlorophyll, a FlowCam LO which has been designed for pharmaceutical purposes and a FlowCam Nano wich is designed to image particles between 300 nm to 2 µm. There is a risk that, in the future, Yokogawa Electric Corporation will invest more into FlowCam applications for life sciences than for environmental sciences.

As a flow cytometer, the FlowCam, can use different types of triggers for the imaging: they can be set to auto-trigger or fluorescence (by photosynthetic pigments). If it is set to fluorescence for particles in the size range of phytoplankton, particles without fluorescence are not imaged and therefore missed. This could be the case of ciliates or smaller rotifers. The auto-trigger relies on scattering signals of the flow cytometer. For larger and more rare plankton, more volume needs to be analysed in order to get a representative sample. Besides the trigger, the flow rate and the number of frames per second have a large effect on the proportion of particles within the sample that are effectively being imaged. Those settings need to be optimised in order to not image the same particle multiple times while imaging the largest possible proportion of the sample. The machine can estimate that proportion based on the flow and the frame rate. At the same time, it has been shown that numbers of 1000 particles per mL should not be exceeded in order to prevent overlapping of particles (Camoying and Yñiguez 2016). For this, the sample density will have to be assessed before measurement with other methods (e.g. Chl-a fluorescence).

In order to image particles with the highest resolution possible, the right magnification needs to be used. For each magnification, a preset matching flow cell and syringe are available (**Fig. 2B**). Each flow cell has its own settings and maximum particle diameter which is given by the shorter side of the flow cell. Therefore, a sample needs to be split by size. The maximum particle diameters for each magnification are the following: 1000 μm (2x), 600 μm or 300 μm (4x), 100 μm (10x), 50 μm (20x). There are two ways of sample preparation (**Fig. 2A**): either serial filtration and re-suspension of each size fraction or serial removal of the largest fraction by filtering, imaging and ignoring all particles smaller than the next smaller fraction. The goal is to image each particle in the highest magnification possible. For the first method, particles might get stuck in the filter/sieve or it might disrupt colonies or single cells. For the second method, the elimination of smaller particles from the analysis relies on the estimation of cell diameter by the FlowCam which might not be as accurate for certain cell shapes (see (Wong et al. 2017)).

**Fig. 2:**
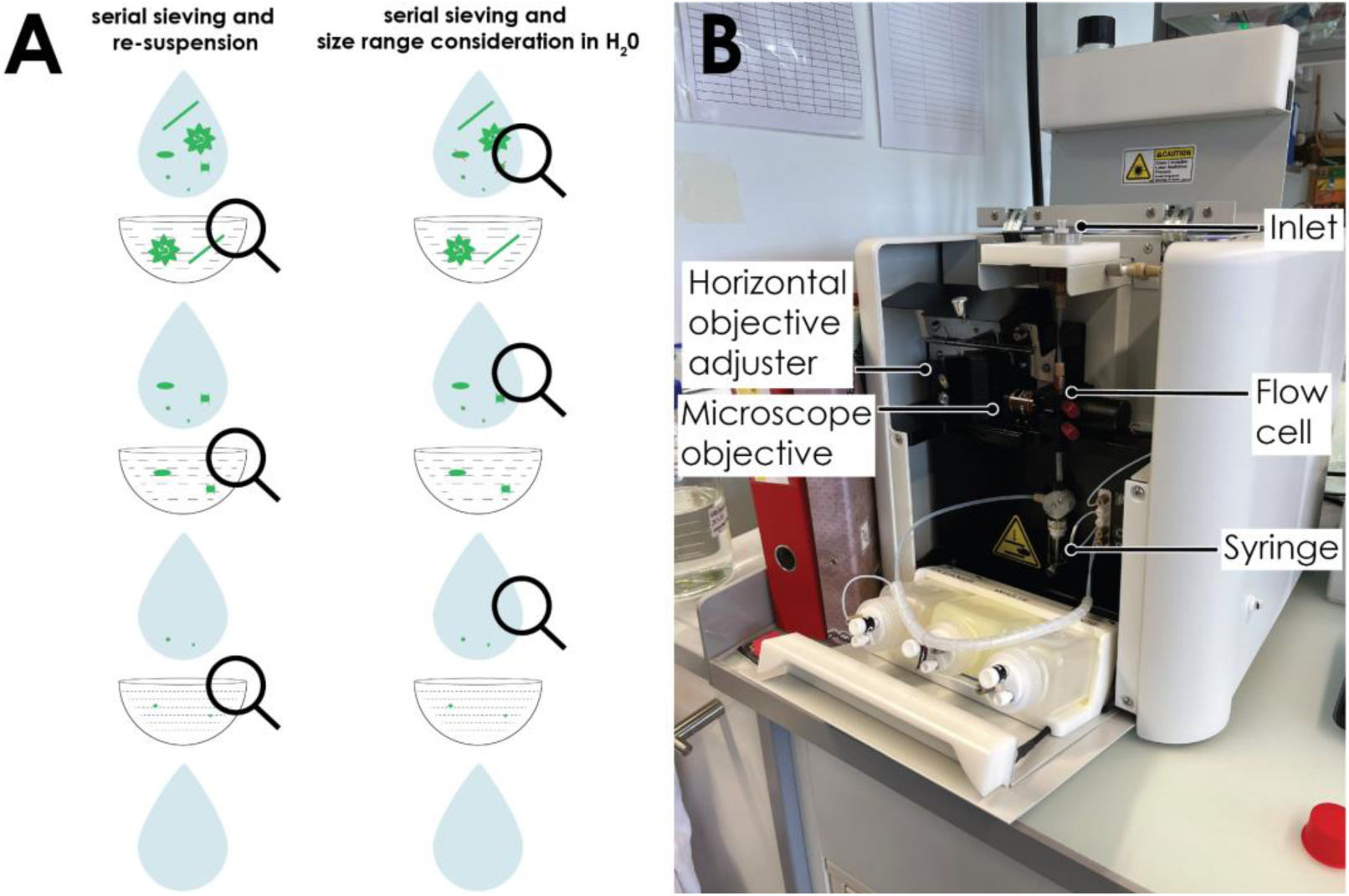
A) Sample processing steps for the use of the FlowCam: either i) serial filtration and re-suspension or ii) serial elimination of larger particles by filtration/sieving and ignoring of smaller particles. **B)** FlowCam structure with three interchangeable parts: objective, flow cell and syringe.

For each lake and instrument, the settings of the FlowCam would need to be adjusted in order to get the sharpest images possible. This is due to differences in sample density, taxonomic composition and hardware. Then, images would need to be annotated into libraries for future automated image classification. For the annotation, FlowCam has a built-in annotation tool that facilitates the annotation and speeds up the process a lot. An alternative would be the use of EcoTaxa (Picheral, Colin, and Irisson, n.d.) for image annotation. Eco Taxa has a built-in random forest classification model for multiple imaging technologies, including FlowCam. One can use the images of other users to speed up their classification. In order to use it, the images from the FlowCam need to be exported and morphometric measurements need to be extracted with a Software called Zooprocess (“Zooprocess : FlowCam & Zooscan” n.d.) that is used with ImageJ. If a custom made classifier was required, an expert in ML would be needed.

#### Aquascope

The Aquascope is designed as an automated *in-situ* monitoring instrument (Merz et al. 2021). It is a dual-magnification microscope camera with a detection range between ∼ 10 μm - 150 μm (5p0x - 5x magnification, pixel size = 0.62 μm) and ∼ 100 μm - 1 cm (0p5x - 0.5x magnification, pixel size = 6.2 μm). The distance between light sources and objective is 5 cm, which allows transmission of light in turbid waters (e.g. during thick algal blooms). The Aquascope images all translucent objects passing between the light source and the objective that are in focus (and this depends on the setting of the image segmentation algorithm) at a desired frequency of up to 10 frames per second. We use an imaging frequency of 1 frame per second and run the Aquascope *in-situ* every hour of the day for 10 minutes.

##### Aquascope lab-adapter

The Aquascope is not directly applicable to measurements in the lab. We developed a lab-adapter that works as a flow-through system. In Fig. 3, you can see the Aquascope in the lab with the lab-adapter. The adapter consists of a PVC-pipe that is installed between the two sides of the DPSC and isolated with a silicone sheet isolation. On the top, a funnel can be inserted into the lab-adapter which functions as an inlet. The chamber holds 0.8 litres while the funnel can hold another litre. The chamber should not run dry while it is taking images which restricts the sample to a minimum of two litres. At the bottom of the chamber, there is an outlet valve which can regulate the flow rate of the sample.

**Fig. 3:**
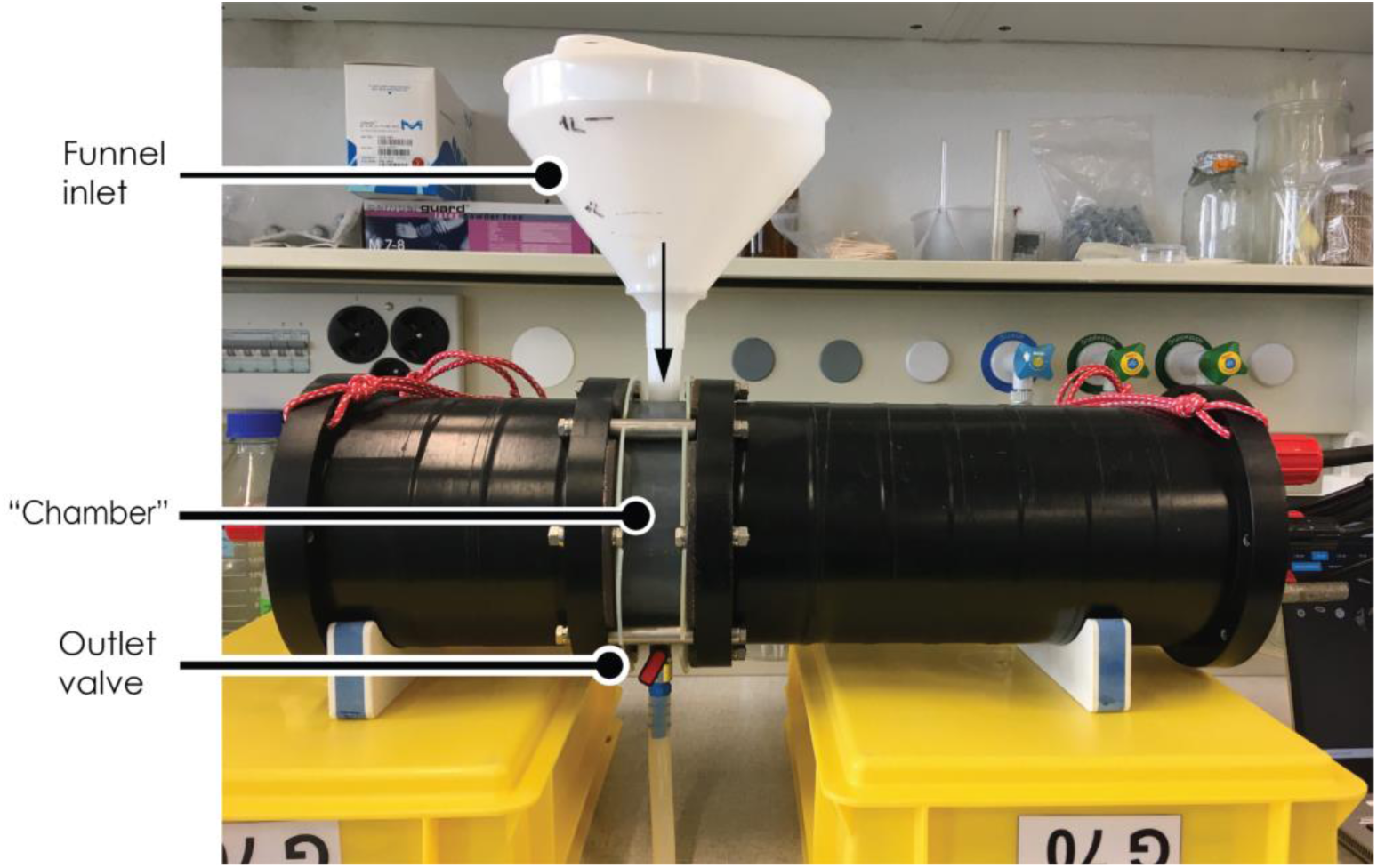
Aquascope lab-adapter. A sample is filled into the chamber through the funnel inlet and flows out through the outlet valve. The flow can be adjusted with the outlet valve.

We have noticed that fast flow rates (> 0.5 L/min) lead to a less consistent flow than slow flow rates. This is most likely the case due to the larger change in water pressure from the top with faster emptying funnel volumes. For flow rates around 0.2 L/min, one gets a constant flow with no risk of an empty chamber within 5 minutes. At this rate, repeated imaging of the same object is limited as can be seen in Fig. 4.

**Fig. 4:**
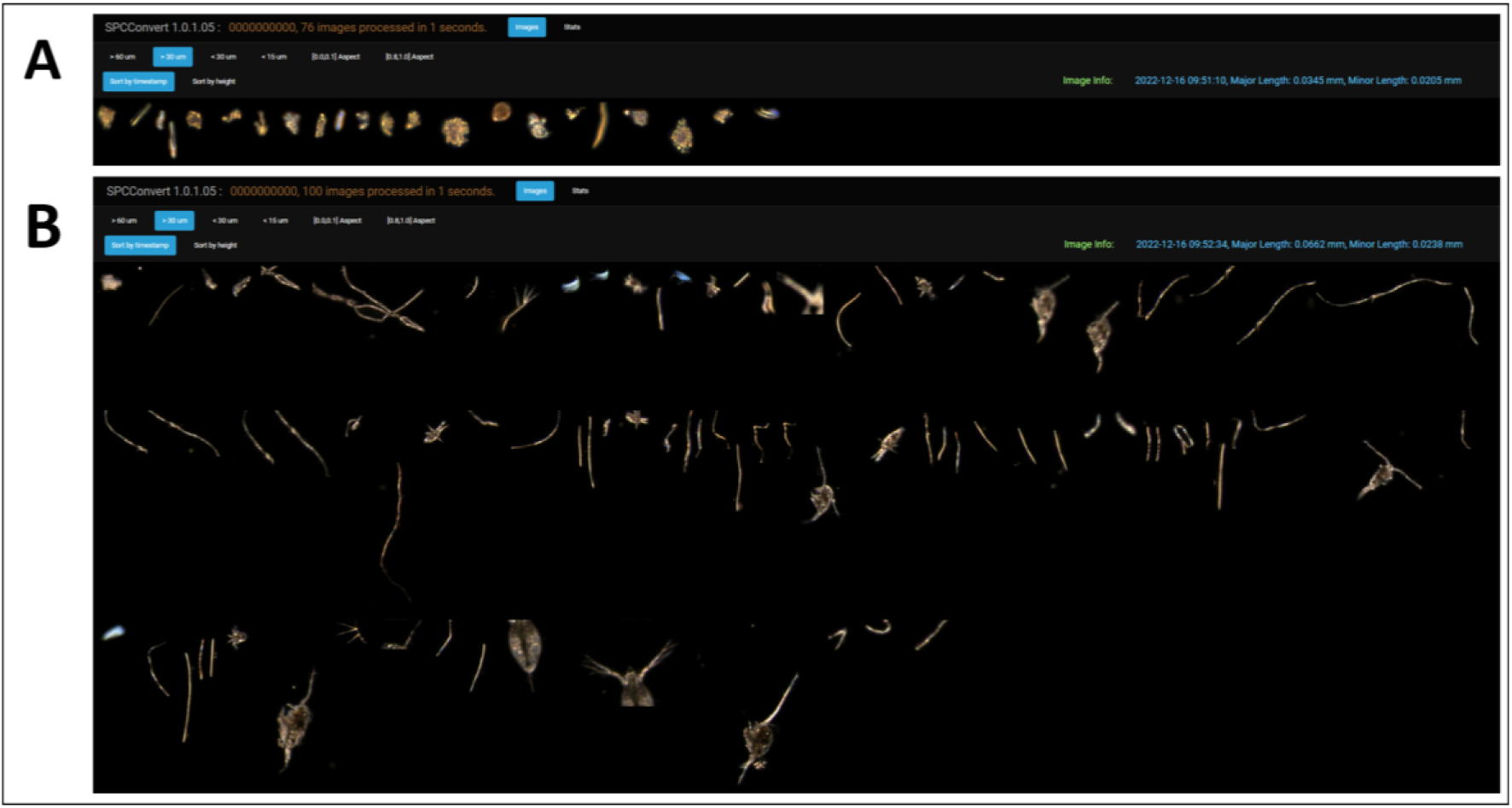
Objects above 30 μm (5p0x) or 300 μm (0p5x) sorted by timestamp in the **A)** 5p0x magnification and **B)** 0p5x magnification. The images were taken from an IWS sample from Murtensee in December 2022 within 5 minutes and a flow rate of 0.25 L/min.

Another factor to consider for the imaging of plankton samples, apart from the flow rate, is the total time the sample is imaged at the time. One needs to take the two following points into consideration when choosing the runtime: a) Plankton starts to sediment in the chamber when the camera runs too long. Therefore, after a while it will take more images than at the beginning. This is especially relevant for slower flow rates. b) Depending on your sample volume and flow rate, you might need to refill the funnel or keep the flow rate slower and/or the runtime shorter. Emptying the chamber in between imaging runs and not refilling the funnel during the run resembles real replicates with a random subset of the entire sample being imaged. Refilling the chamber with the same sample in the same run is not encouraged.

Further minor things to consider are not to start the camera while filling the chamber. Bubbles will be imaged which will leave you with many large images of bubbles. Additionally, we recommend emptying the camera between runs to create real replicates. If measuring multiple lakes or samples in a row, we recommend flushing the chamber, funnel and outflow tube with clean (e.g. tap or deionised) water in between samples.

For oligotrophic lakes, such as lake Thun or lake Brienz, we recommend using a net sample. For lake Thun (09.08.2022), within 5 minutes we got on average 56 images in the 5p0x magnification and 89 images in the 0p5x magnification. For lake Brienz (13.09.2022) within 5 minutes we got on average 92 images in the 5p0x magnification and 52 images in the 0p5x magnification. In comparison, lake Hallwil yielded 230 (24.08.2022) and 173 images (21.09.2022) in the 5p0x magnification and 152 (24.08.2022) and 302 images (21.09.2022) in the 0p5x magnification. If one’s goal is to image a robust estimate of the lake’s species composition, using a net sample is much more efficient.

##### Data processing and analysis

After imaging your samples, the data is saved in a tar-file, which is always called 0000000000.tar. A tar-file (Tape Archive file) is an archive file generated by a linux computer that contains multiple files (e.g. images) and is usually used for storage or sharing. Tar-files do not compress files compared to zip-files. The 0000000000.tar file needs to be extracted and the black-and-white images inside that folder (the full frames are not saved) are then converted into colour images (**Fig. 5**; for a comparison with FlowCam images see **Fig. S3**). At the same time, features (e.g. area, major axis length, solidity) are extracted from the image and saved into a features.tsv file using a python script called SPCConvert which is an open-source script available on github (Roberts n.d.; Orenstein et al. 2020). SPCConvert is easy to apply using a Linux subsystem on Windows called Ubuntu, which is also free of charge to use. We found that for very large images (e.g. large *Planktothrix rubescens* filaments from Hallwilersee), a normal desktop computer or laptop (16.0 GB RAM; 11th Gen Intel(R) Core(TM) i5-1135G7 @ 2.40GHz processor; 64-bit operating system, x64-based processor) is not powerful enough to convert the images into colour images and a file with features with SPCConvert. The python script runs smoothly on a stronger server such as the one the Research Data Management Project at Eawag owns. Therefore, a requirement for the use and handling of data produced by the Aquascope is access to a server.

**Fig. 5:**
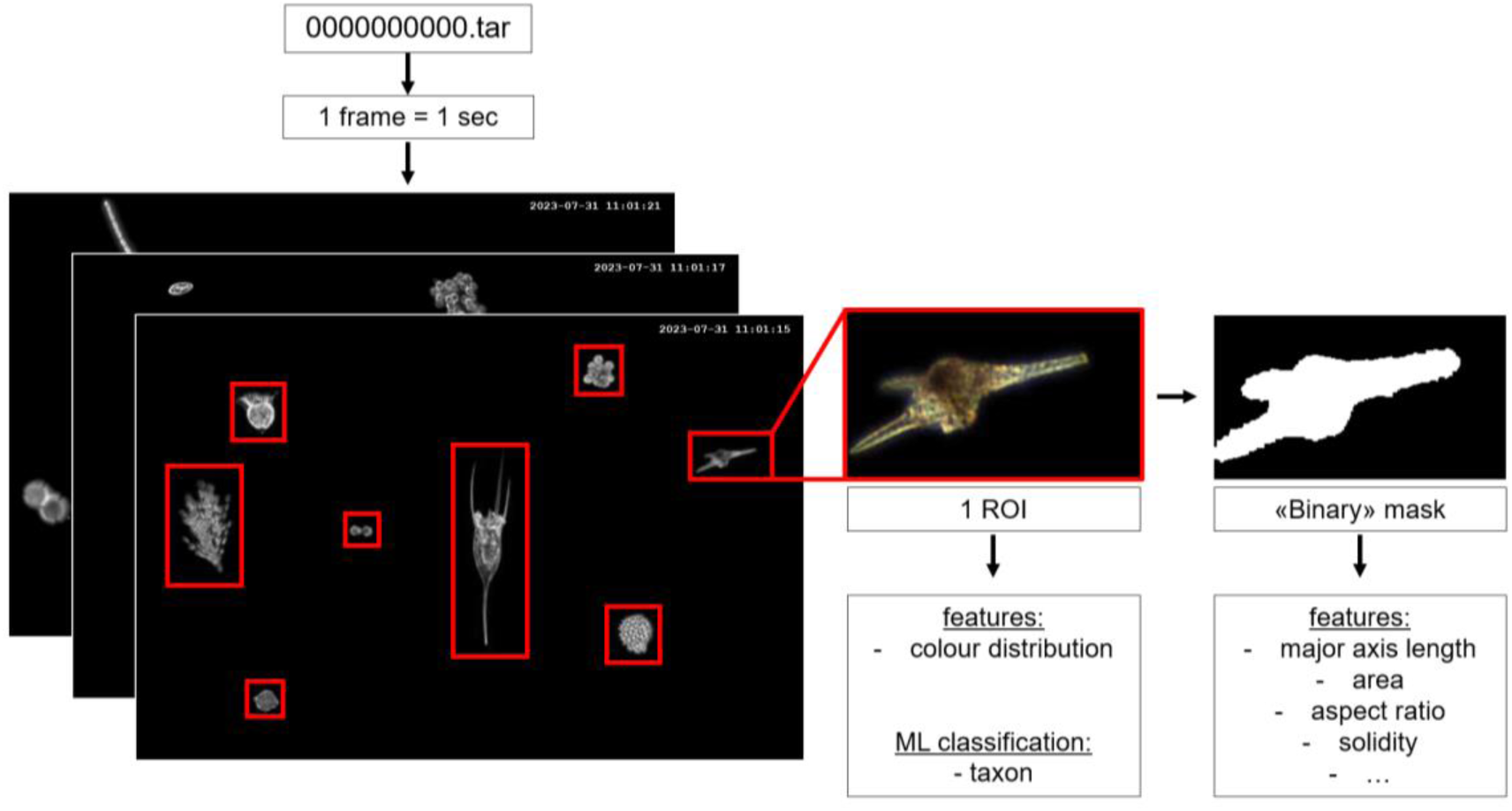
Data processing from raw Aquascope data to features and classification. Each ROI (i.e.cell/colony/individual) is cut from a frame and converted into colour with SPCConvert. At the same time, features are extracted from each ROI. In addition, a ML classifier is applied to the colour images to determine the taxon.

After that, the colour images can be used in a ML classifier which is also based on a python script that is open-source and available on github (Kyathanahally n.d.). The ML classifier currently available is trained on data from the Aquascope installed in Greifensee. The image library used to train the ML classifier was built manually. As of now, there is no built in or compatible tagging software that can speed up the annotation process like it is available for FlowCam (internal annotation or EcoTaxa).

Each Aquascope camera has slightly different hardware and settings which changes the overall colour appearance of the images. In **Fig. 6**, we can see the slightly different colour in the images from the Aquascope we have in the lake, which was used to train the ML classifier, and the images produced in the lab by our second Aquascope. To overcome this difference, there is an open-source python script on github (Kyathanahally n.d.) that is able to transfer the colour of the original DPSC images to the images of another Aquascope.

**Fig. 6:**
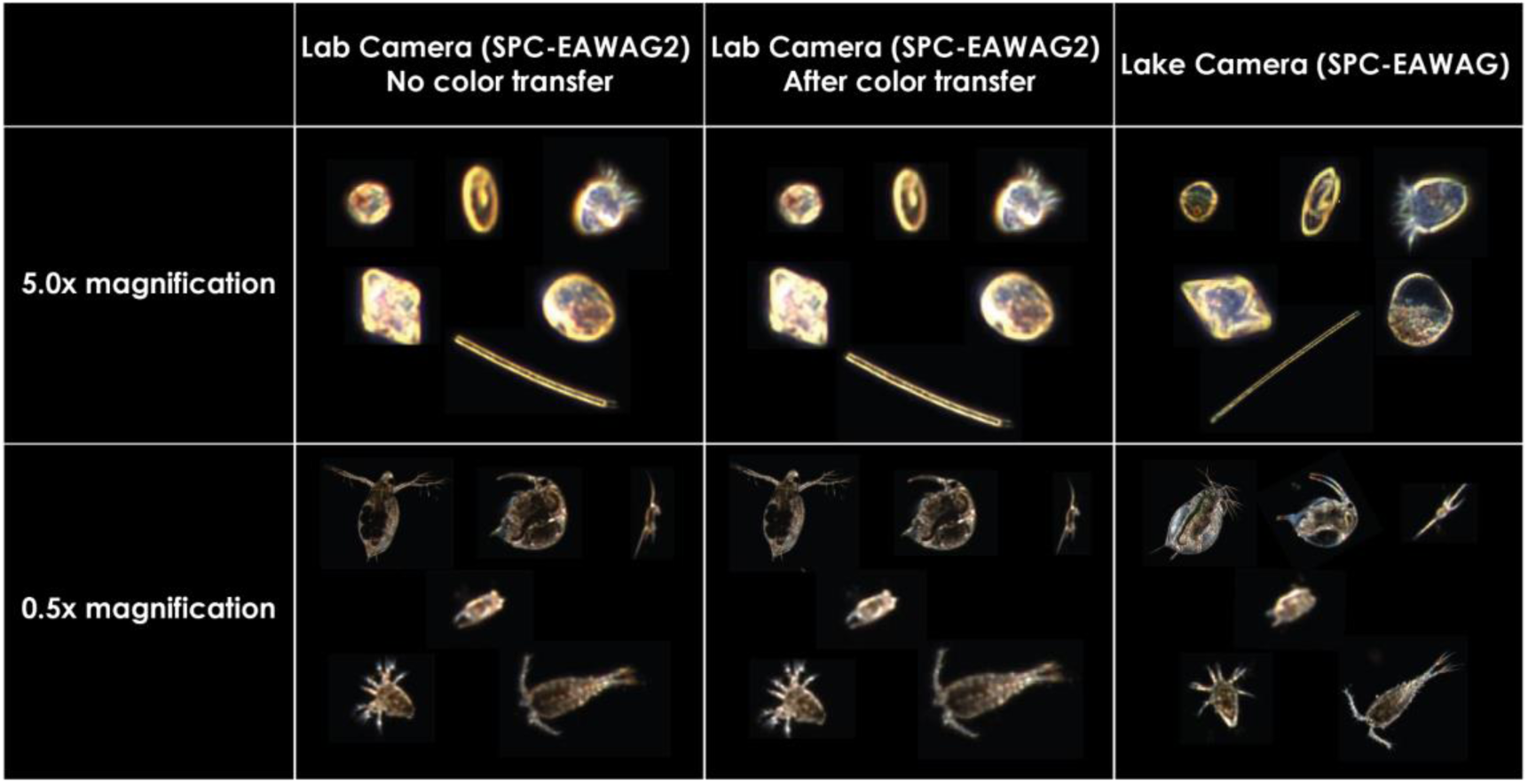
The two Aquascope cameras owned by the Pomati lab in comparison regarding the colour of images they generate. The SPC-EAWAG camera is installed in lake Greifen at 3m depth and the ML classifier is based on those images. The SPC-EAWAG2 is in the lab and used for other field and lab measurements. The lab-adapter was tested with this camera.

All the scripts and softwares described above are free of charge, open-source and easy to use. User-friendly manuals for each step of the process are available.

### Goals

In order to ensure plankton monitoring in the future, potentially with automated imaging techniques, the goals of this projects are the following:

1. Highlight the automated approach’s gaps and opportunities for routine plankton counting and identification. This is accomplished through i) testing the accuracy of classification under laboratory conditions and ii) comparison of the Aquascope to traditional monitoring.
2. Provide the basis for a bigger follow up proposal for in depth benchmarking of automated plankton monitoring techniques and potentially a plankton monitoring centre (i.e. a service and advisory centre for plankton monitoring, counting, and classification).

## Methods

Across the year 2022, we imaged multiple samples from the cantons of Aargau (Hallwilsersee) and Bern (multiple lakes) in the lab. We used the lab camera, with the flow adapter for that. All samples from Aargau were from an integrated tube (garden hose) sample. The samples from Bern were either integrated (IWS) or net samples for phytoplankton and net samples for zooplankton. We resuspended net samples into tap water for analysis and noted the resuspension volume down. If samples from any lake were too dense, we diluted the samples again with tap water and noted the dilution down. We also imaged two samples from Greifensee. One was an integrated (Schröder) sample from the deepest point of the lake (12.04.2023, i.e. the same as traditional monitoring) and the other one was a discrete water (Niskin) sample from 3m depth where the Aquascope is situated in the lake (23.03.2023).

We then imaged the samples with a frame rate of one frame per second. The flow rate was usually in the order of magnitude of 0.2 L/min. We mostly imaged the samples for 5 minutes, then emptied the entire chamber and refilled it for the next replicate.

Afterwards, we extracted each ROI’s area (mm^2^) with SPCConvert and classified each image with our ML classifier. We then averaged the relative abundances of all the taxa we found across all replicated camera runs. Further, we corrected the relative abundances by the dilution factor and the different resuspension volumes where applicable. To generate relative biomasses, we multiplied the abundances with the area of each image.

## Results and Discussion

We provide a large colour legend for all following plots in the supplementary information. Please consult **Fig. S4** for colour information on the taxonomy of each plot.

### Flow rate and density on abundance estimation of Aquascope

In order to understand if the flow rate has an influence on the number of images captured by the Aquascope and how that scaled with the sample density, we performed a series of lab experiments with a variety of dilutions and flow rates. The ROI/sec we get for different dilutions is consistent with our expectations of a perfect dilution series (**Fig. 7A**, **Fig. 7C**). This is true for both magnifications. We only find a significantly different mean ROI/sec to the expected value in the 0p5x magnification at the 1:7 (12.5%) dilution (**Tab. S1**).

**Fig. 7:**
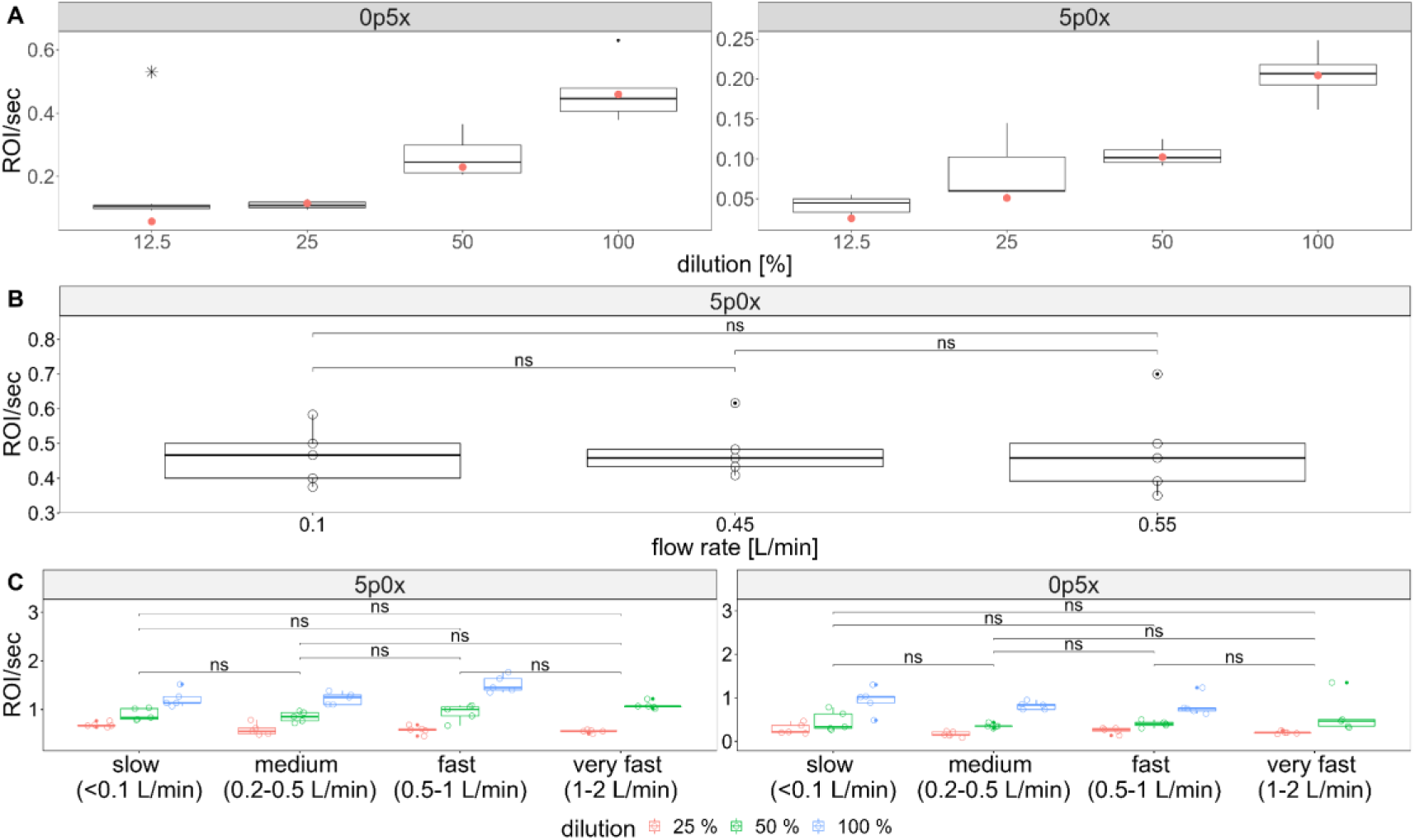
Flow rate and dilution trials with the lab adapter on the Aquascope. Non-siginifcant (ns) T-tests between flow rates are displayed as horizontal bars (panels **B** and **C**). **A)** Dilution series of a sample from Hallwilersee (November 2022). ROI/sec for each magnification are displayed for the dilution of the sample to 50% (1:1), 25% (1:3) and 12.5% (1:7). Red points are expected mean values for a perfect dilution series. The star represents a statistically significant (p < 0.05) difference between the observed and expected mean ROI/sec. **B)** A mixed sample of lab cultures (Oocystis sp. + Lagerheimia sp.) under different flow rates. **C)** A combination of different flow rates and dilutions was tested on a sample from Greifensee (July 2022). The sample was diluted to 50% (1:1) and 25% (1:3).

If we compare flow rates, we find that there are no significant differences in the number of ROI taken per second (**Fig. 7B**, **Fig. 7C**) under different flow rates. Considering this and personal experience that the faster the flow rate, the more inconsistent it is, we suggest using flow rates at the lower end. Under flow rates of around 0.2 L/min, we do not get a large number of objects that get imaged repeatedly (**Fig. 4**) and images are considerably sharp which improves classification.

### Accuracy of the classifier in the lab application of Aquascope

To understand how well our existing ML classifier, that is based on the Aquascope installed in Greifensee, performs under laboratory conditions with another machine, we imaged an integrated Schröder sample from Greifensee with the Aquascope in the lab. The sample was taken in the same way as the monthly monitoring that contributes to the long term monitoring from Canton Zurich. We ran the 2 L sample 10 times through the lab adapter for 5 minutes. This resulted in a total of 1270 ROIs in the 5p0x magnification and 3433 ROIs in the 0p5x magnification. We then classified each image before and after colour transfer. We manually classified all images in order to be able to calculate the accuracy of the classifier. We find that the colour transfer does not improve the accuracy of the classifier (**Fig. 8, Tab. S2, Tab. S3**). The number of ROIs that are used as a reference for the colour transfer does not seem to have a strong effect on the accuracy of the classifier. In our test, we only found a slight negative effect with increased reference sample size (**Tab. S2**). Even though the colour transfer does not seem to improve the overall classification, for some taxa there is an improvement in the classification accuracy (**Fig. 8, Tab. S3**). The largest improvement in classification accuracy is in the classes rhodomonas (from 0.75 to 0.9167/1) and cryptomonas_cryptophyceae (from 0.7273 to 0.8182). For the rotifer classes trichocerca and polyarthra, the colour transfer decreases the accuracy of the classifier (trichocerca: from 1 to 0.0952/0.0476 and polyarthra: from 0.7059 to 0.3922/0.3333).

**Fig. 8:**
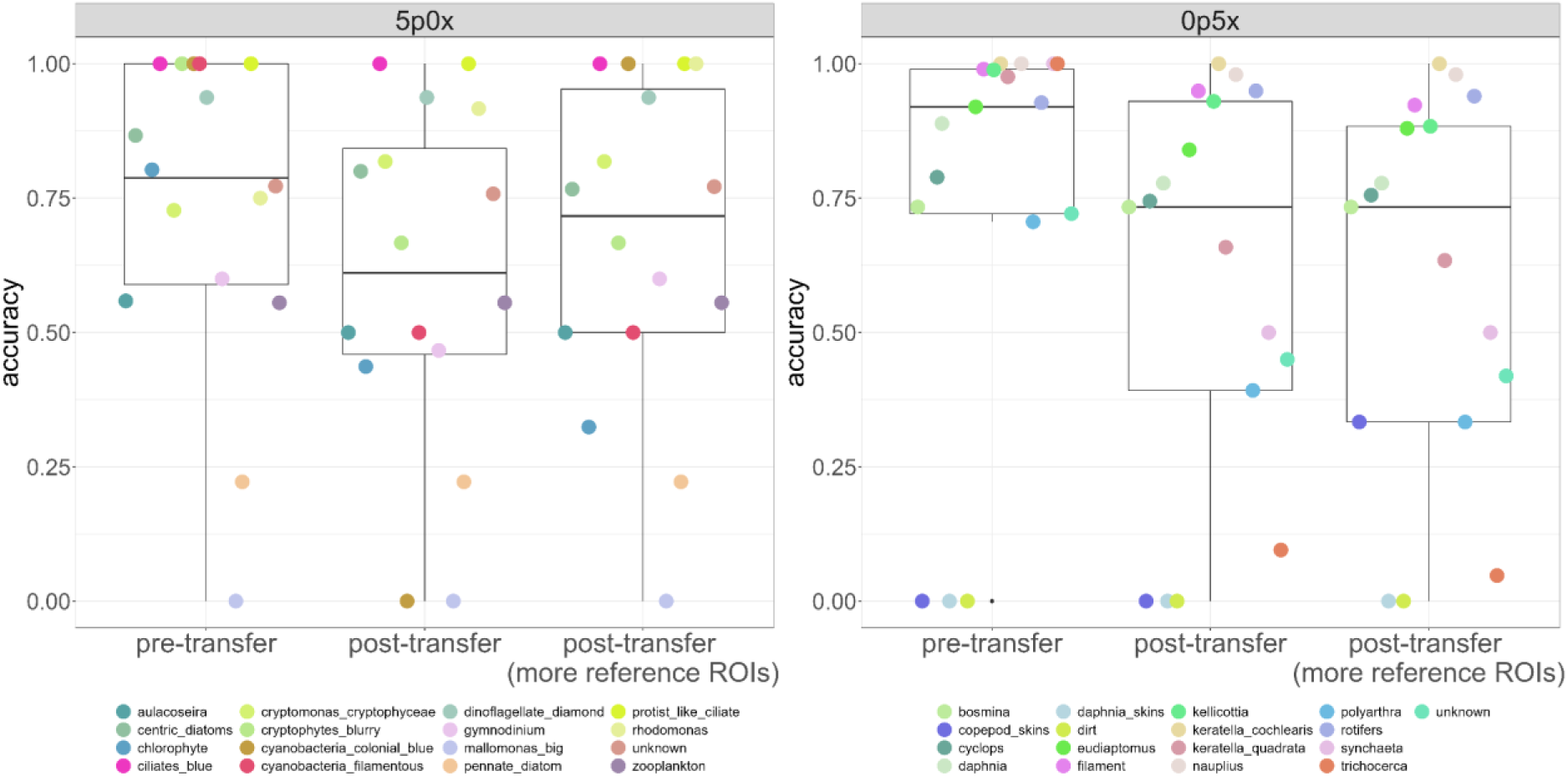
Accuracy of the classifier on a sample from Greifensee (12.04.2023), imaged in the lab. The accuracy of each taxon of the 0p5x and the 5p0x magnification was assessed before and after using the colour transfer script. For the colour transfer, we used 1727 ROIs or 5178 ROIs from the 0p5x magnification and 6219 ROIs or 13287 ROIs from the 5p0x magnification as a reference. Each point represents one taxon and its accuracy in the three different colour transfer levels (x-axis). The number of images per taxon can be found in **Tab. S3**.

From validation and testing of the ML classifier (Greifensee lake camera images only), we know that the classifier has an accuracy of 90.3% on classifying images from the 5p0x magnification and 93.8% with images from the 0p5x magnification (Kyathanahally n.d.). Therefore, we can see that using a second camera in the lab, we lose an accuracy of about 13.6% in the 5p0x magnification and 2.3% in the 0p5x magnification (**Tab. S2**).

As we can see in **Fig. 9**, this could be due to the fact that we have a larger proportion of unknowns when we run the camera in the lab with a sample from the lake (Niskin bottle from 3m; 5p0x: 49.70 %; 0p5x: 44.47 %), compared to the data generated by the camera in the lake (5p0x: 23.80 %; 0p5x: 9.94 %). This is more pronounced in the 0p5x magnification.

**Fig. 9:**
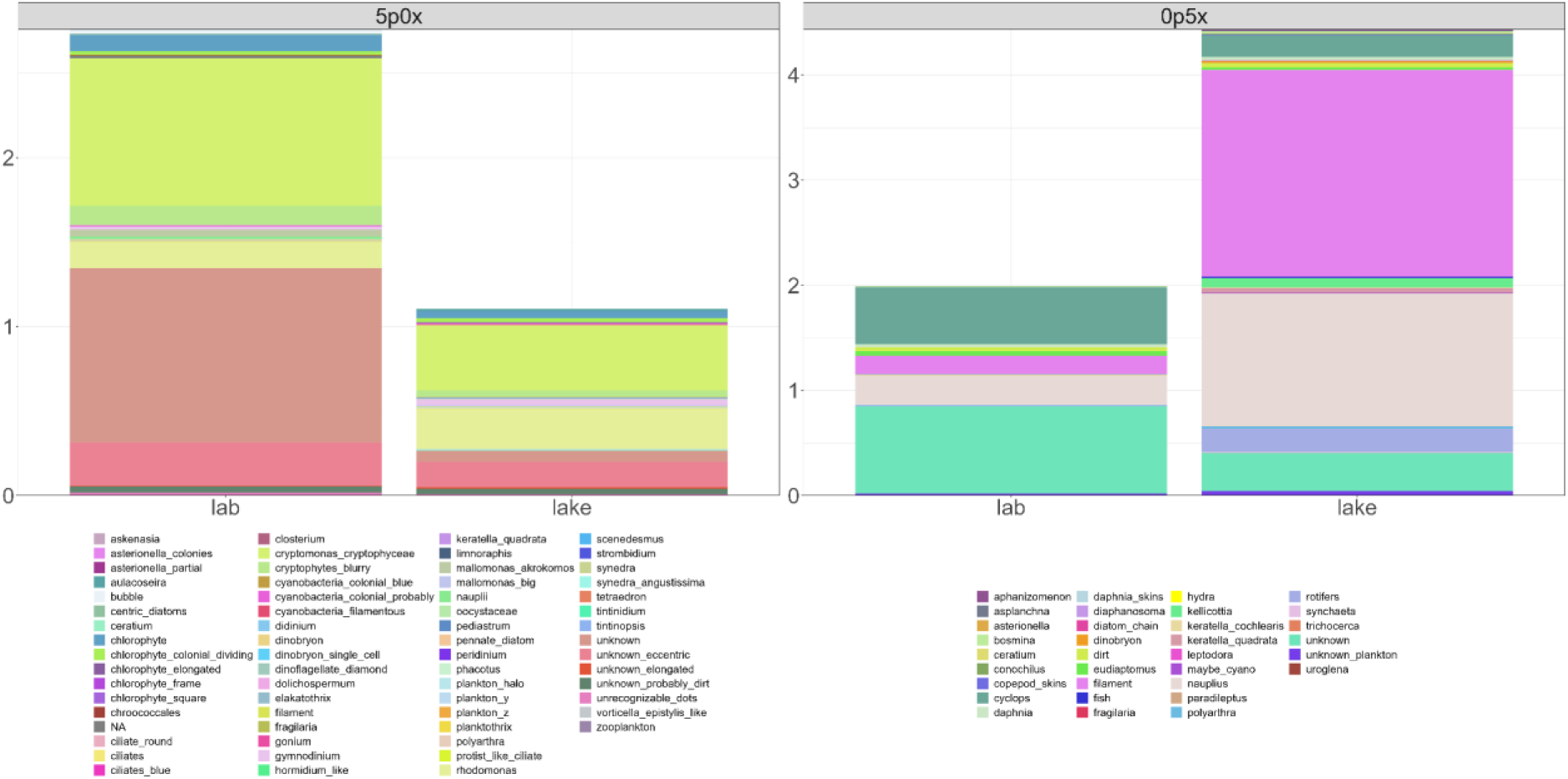
Comparison between a 3m sample from Greifensee (Niskin, 23.03.2023, “lab” column) and the images taken at 3m with the lake camera (“lake” column) on the same day (23.03..2023 00:00 - 23:59). Separated by magnification into two side-by-side plots (left: 5p0x magnification, right: 0p5x magnification). Displayed are the mean ROI/sec across all lab camera runs and the ROI/sec from the lake camera.

This might be due to more blurry images caused by the flow and turbulence in the lab adapter, increased dirt and dust particles that fall into the sample while running it or disrupted cells, colonies or individuals due to the handling of the sample. If we remove the unknown classes, we see that the match of the relative abundance of the classes improves (**Fig. S5** and linear model fit for i) 5p0x magnification: slope = 1.83, p <2e-16, Adjusted R^2^: 0.8526 ii) 0p5x magnification: slope = 0.14, p = 0.0047, Adjusted R^2^: 0.2253).

In order to capture a representable sample from the lake community with the Aquascope, we need to increase the sampling effort, either by running the sample longer in each replicate or having more replicates. We can see in the taxon accumulation curves in **Fig. 10** that the number of taxonomic groups detected in the 0p5x magnification does not change after 5 replicates of 5 minutes each (1657 ROI). The 5p0x magnification on the other hand benefits from a larger amount of replicates, especially since the image volume is much larger than in the 0p5x magnification. Within 10 replicates (1270 ROI), we do not see a saturation of taxonomic groups detected as we see in the 0p5x magnification.

**Fig. 10:**
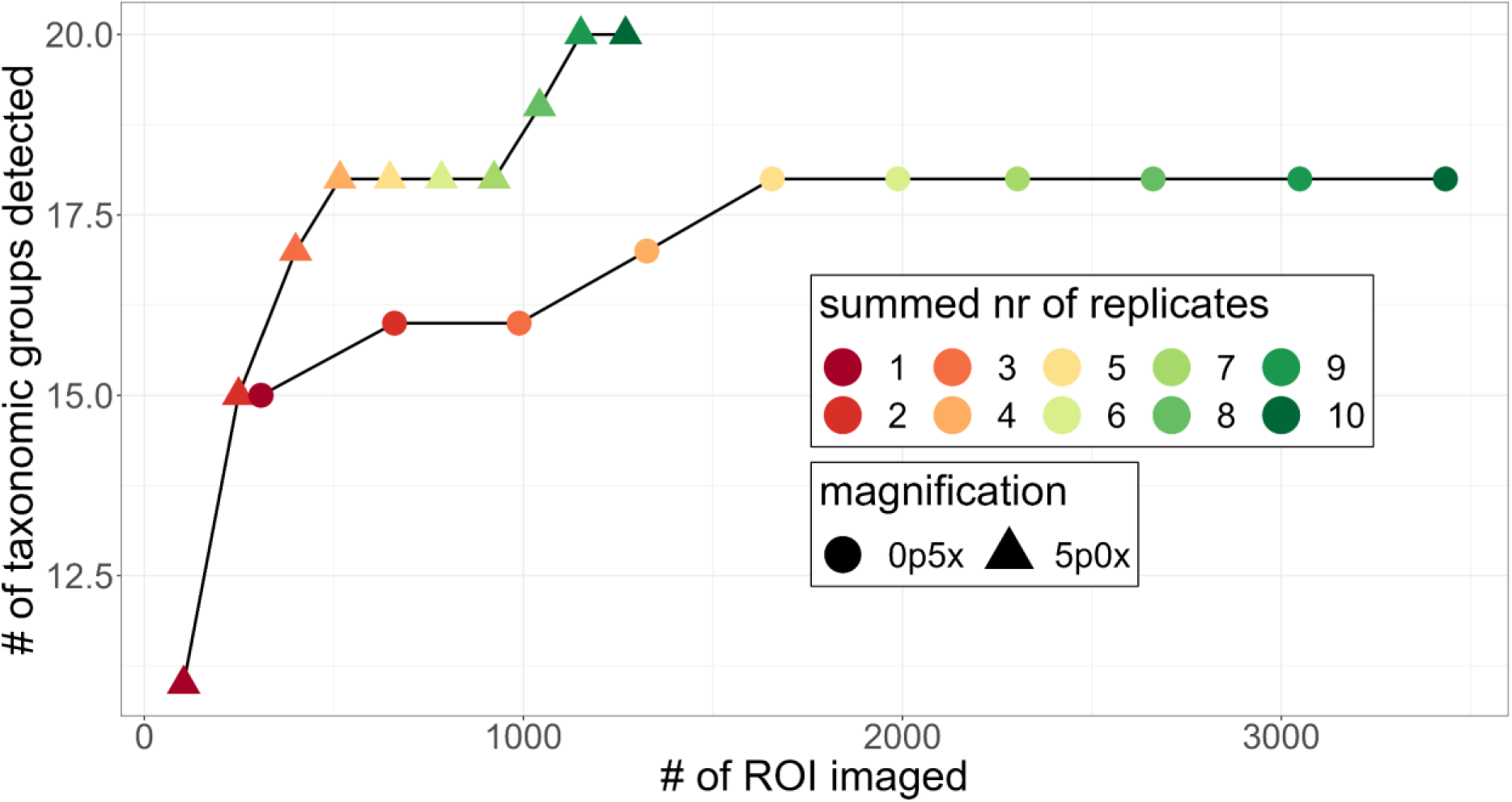
Taxon accumulation curves for an integrated sample from Greifensee (Schröder 0-20m, 12.04.2023) measured with the Aquascope in 10 replicates of 5 minutes each (1 frame per second). One line and shape for each magnification and the cumulative number of replicates in colour.

Ultimately, we are interested in a representative sample of all taxonomic groups in the sample that have a known classification. Due to the higher proportion of unknown images produced in the lab (**Fig. 9**) and the results from the taxon accumulation curves (**Fig. 10**), we recommend taking a larger amount of images than required by the minimum number of objects in order to have a robust estimate of the plankton community composition in the sample. In addition, we think that it is necessary to adapt the ML classifier for images taken in the lab as they are not only different in colour due to hardware differences, but also in the quality of the images which might be biasing the classification. In addition, for different lakes, there might be taxa that are not included in the ML classifier for Greifensee. Those taxa would have to be added to the classifier.

### Quantification of Plankton and comparison to microscopy counts

To understand how well the Aquascope reflects the traditional monitoring, we compare samples run in the lab with traditional microscopy data. We did this for multiple lakes. We compared different levels of data treatments with the microscopy data, i.e. i) raw ROI/sec ii) ROI/sec adjusted for the imaging volume differences between the two magnifications and iii) calculated cells/mL based on lab cultures (Merz et al. 2021). In the following, we will discuss the different approaches, how well they reflect microscopy data.

The two magnifications have different theoretical volumes that are being imaged. Those differ by up to a factor of 1000 (5p0x = 0.2 - 10 μL, 0p5x = 4 - 200 μL). According to our analysis, the 0p5x magnification detected 23 times more *Asterionella* ROIs than the 5p0x magnification (**Fig. 11**). We therefore divided the ROI/sec from the 0p5x magnification by 23 before summing across magnification.

**Fig. 11:**
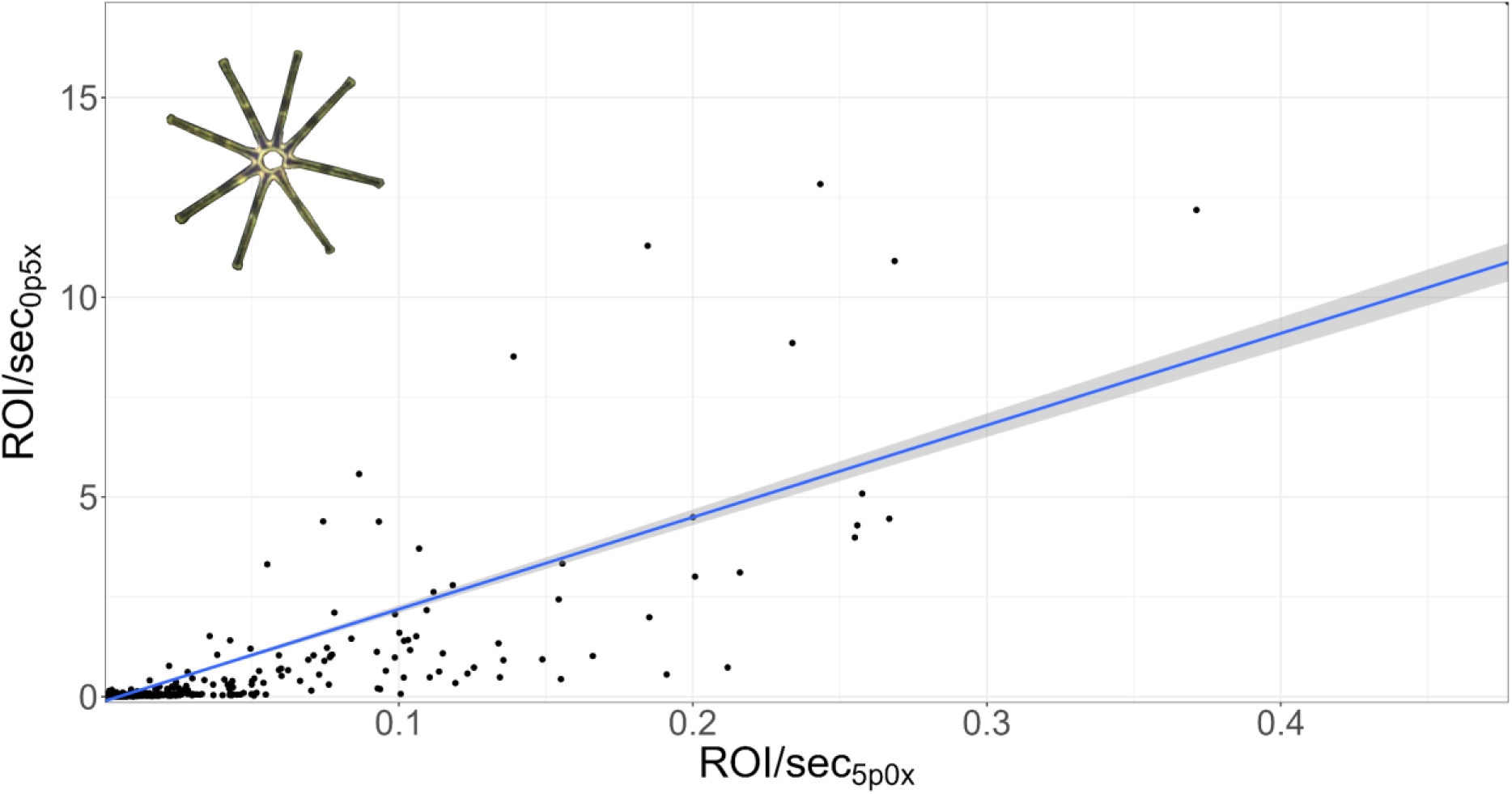
Comparison of ROI/sec from both magnifications on Asterionella sp. Asterionella sp. has a colony diameter of about 100 μm which lies well within the detection range of both magnifications. The slope of the linear relationship between the two magnifications is 23 with an intercept of −0.11 (R^2^ = 0.659).

To convert ROI/sec to cells/mL, one can use the calibration in Fig. 3 in Merz et al. (2021). Then, we should be able to sum abundances across the two magnifications. If we then want to calculate biomasses, we can correct the total area of each taxon per day by the correct abundance in cells/mL.

We found that our best fit between microscopy data and data from the Aquascope is between biomass/L (microscopy, relative abundance on phyla level) and the bioarea/L based on the total bioarea and the cells/mL (Merz et al. (2021) calibration). A linear model between the relative biomass of the different phyla gives a slope of 0.65 with an adjusted R^2^ of 0.37 (see other fits in **Tab. 1**). The imperfect linear model fit could be due to either an inaccurate estimate of abundance and thus total bioarea, or due to misclassification in Aquascope. Alternatively, it can be due to inaccuracies in the calculation of biomasses from microscopy counts, where a nominal cell size is assigned to each taxon, sacrificing individual level variation.

**Tab. 1:**
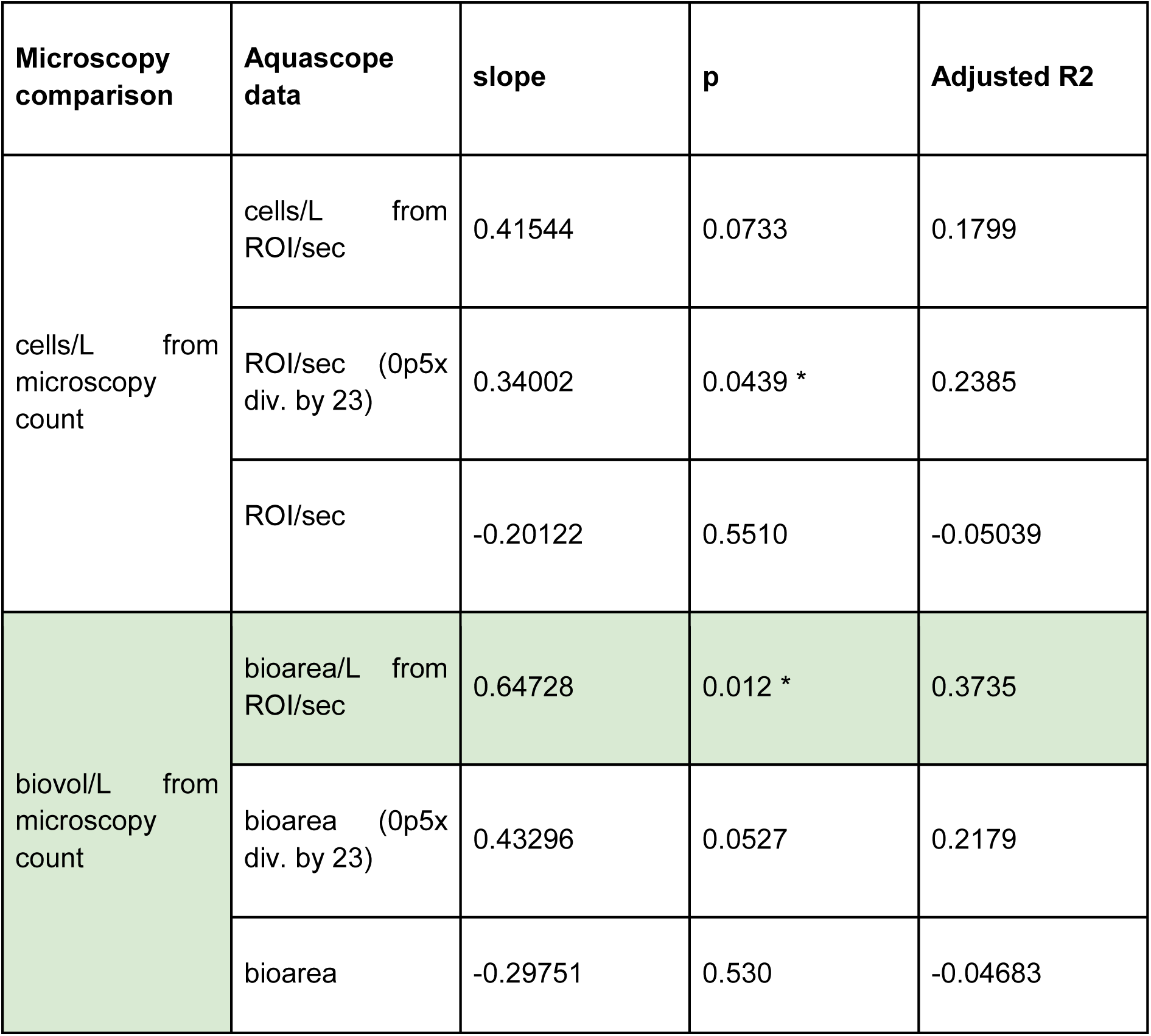
Relative abundance (%) of each phylum from an integrated sample from Greifensee (Schröder 0-20m, 12.04.2023) measured with traditional microscopy and the Aquascope aggregated into phyla (see Fig. 12). Linear model statistics where the Aquascope data was fitted against the microscopy comparison (lm(Aquascope data ∼ microscopy comparison). The best fit is highlighted in green.

We further found that the conversion from ROI/sec to cells/mL does not work well for colonial algae because with the Aquascope, one ROI represents one colony or filament and with traditional microscopy, each cell is counted as one unit. In addition, the calibration of phytoplankton abundances from Merz et al. (2021) does not include large colonies and is based on laboratory cultures. An extended calibration with larger colonies and more species might increase the fit between microscopy counts and the abundance from the Aquascope, especially for colonial plankton. As an example, cyanobacteria dominate the microscopy counts from Bern while they are much less represented in the Aquascope lab data (**Fig. S7, Fig. S8**). Note that we assume that the classification of the images is 100% correct. We know that this is not realistic at this stage (**Tab. S2, Tab. S3**) but it gives us a good indication on the composition of the plankton community in the lakes of the canton of Bern. When we compare the bioarea composition from the Aquascope and the biomass data from microscopy (**Fig. S7, Fig. S8**), we see a more similar composition that is not as strongly biased by Cyanobacteria anymore. This is most likely due to the reason mentioned above where the abundance estimate of colonial plankton is different in traditional microscopy and the Aquascope.

In addition, we can see that with the Aquascope, we can enumerate microzooplankton (Rotifera and Ciliophora) which are not included in the traditional monitoring campaigns of some cantons (e.g. canton Bern or lake Zurich (WVZ), **Fig. S7**). Microzooplankton has been shown to be an important link in the aquatic food web (Merz et al. 2023) and we therefore recommend including them in the enumeration of plankton samples.

In Hallwilersee, we only have biomass estimates from traditional microscopy. Hans-Ruedi Bürgi delivers data for phytoplankton and microzooplankton in g NG/m^2^ for an integrated sample and for crustacea from a net tow. Due to the difference in sampling method for the crustacea (net sample in microscopy and integrated sample in the Aquascope), the bioarea from the Aquascope and the biomass from the microscopy counts are largely different (**Fig. S9**). Excluding the crustacea from the comparison, we find that the biomass from microscopy on phyla level does not reflect the bioarea from the Aquascope (**Fig. S10**, linear model fit: slope = 0.0926, p = 0.023, Adjusted R^2^: 0.0147). This might partially be because *Planktothrix rubescens* is seasonally highly abundant in Hallwilersee. With the camera, we do not have a category for that species in the 0p5x magnification. It often lands in the category filaments which we cannot distinguish from other filaments with the ML classifier. That class made up a large proportion of the classified images in multiple samples (**Fig. S11**). We also found a large proportion of unknown images (note that we might be misclassifying images at this stage). Therefore, those images cannot be assigned to a phylum and bias the comparison of the microscopy and camera data. This shows the need for an extensive adaptation of the ML classifier to cover multiple lakes with different species compositions. With an adapted ML classifier, we expect an improved fit between microscopy and camera data and therefore the possibility to continue the long term time series on a broad taxonomic level.

In Greifensee, where we manually checked the classification (Schröder 0-20m, 12.04.2023), we can see that the camera underestimates the relative abundance (in ROI/sec and cells/mL) and biomass of cryptophyta and ochrophyta (**Fig. S6, Fig. 12**), within the latter we are only able to distinguish a total of 7 taxa with the Aquascope (one present in the sample) compared to the 17 ochrophyta taxa regularly identified by microscopy counts (7 taxa in the sample). This might be due to a size constraint posed by the Aquascope. At the same time, we overestimate, relative to microscopy counts, the relative abundance of bacillariophyta, especially in the biomass estimate, and chlorophyta. When we transfer abundances into biomass, we can see that the relative biomass of the phyla chlorophyta and dinophyta in the Aquascope match the biomasses of those phyla in the microscopy counts a lot better (**Fig. 12**).

**Fig. 12:**
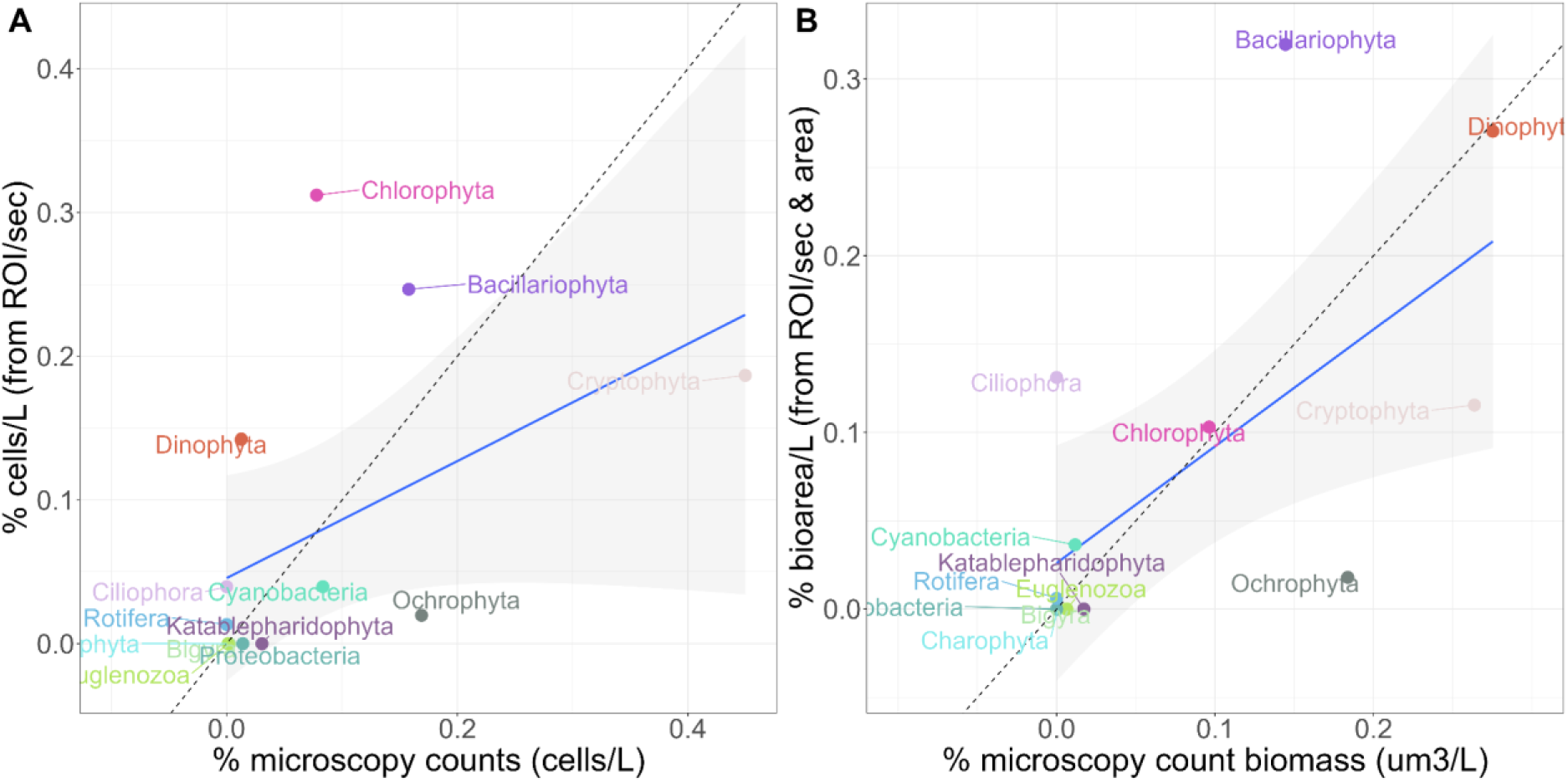
Relative abundance (%) of each phylum from an integrated sample from Greifensee (Schröder 0-20m, 12.04.2023) measured with traditional microscopy and the Aquascope aggregated into phyla (coloured points). The dashed line is the 1:1 line. The blue line with the grey area is the linear model across all points with its 95%-CI (for the linear model, see ***Tab. 1***). Relative abundance of **A)** abundance: cells/L from microscopy counts (x-axis) vs. cells/L calculated from ROI/sec (y-axis). **B)** biomass: biovolume/L (μm^3^/L) from microscopy counts (x-axis) vs. bioarea/L (mm^2^/L) calculated from the total bioarea and the cells/L.

As automated imaging techniques can estimate the two main dimensions of the body size of each individual, biomass (in this case bioarea) estimates might be more accurate than using microscopy counts with nominal biomasses assigned to taxa, regardless of individual level variation (which could be significant, especially for organisms that have juvenile stages or colonies). At the same time, with automated imaging techniques, it is not necessary to use fixatives to count plankton. It has been shown that fixatives bias the size, shape and general appearance of plankton which could lead to inaccurate biomass estimates (Lombard et al. 2019). Nevertheless, biomass estimations are only possible in 2D from images (mm^2^). In theory, SPCConvert reports an estimated biovolume from the shape in mm^3^ but that estimate is based on the assumption that the 3rd dimension is equal to the second linear dimension detected (i.e. objects are deep as they are wide). Therefore, we have not used that estimate and only relied on the biomass estimation in two dimensions. That estimate has been shown to be comparable to microscopy measurements (Merz et al. 2021). In traditional monitoring, standard biomasses are either in g or in μm^3^. Therefore, the difference in the unit used to estimate biomass might further complicate the comparison between total biomass from microscopy and the Aquascope. In the future, a taxon-specific calculation of biovolume (mm^3^) from bioarea (mm^2^) could improve that relationship.

It is difficult to disentangle the factors influencing the differences in relative abundance of taxonomic groups between traditional monitoring and the use of the Aquascope, as each approach has its own intrinsic biases. Some confounding factors include: stochasticity in the sampling of the lake community, stochasticity in the subsampling which is used for microscopic (and Aquascope) enumeration, differences and uncertainties in biomass assignments to counted individuals, and many more. Therefore, extensive benchmarking is needed to match traditional monitoring data and Aquascope data. Likely, the outcomes of further comparison will highlight that each approach has advantages over the other, and inherent limitations. Whether a method should be preferred over the other, really depends on the study objectives and requirements.

### Taxonomic resolution

As we have already seen in the previous sections, the match between the relative abundance of microscopy counts and the one generated by the Aquascope is not optimal. In **Tab. 1**, we saw that the best fit we get between the two is between the biovolume from microscopy and the estimated (by calibration) bioarea of each phylum. We have also seen that we can distinguish less taxa with the Aquascope than with traditional microscopy (e.g. ochrophyta). If we now compare the loss of data at a finer taxonomic level (**Tab. 2**), we can see that Aquascope data lose more information a fine taxonomic levels compared to the microscopy data, which is counted at a higher taxonomic resolution (e.g. 60.76% loss at the order level in the Aquascope data and 12.54% in the microscopy data). At the same time, the linear fit between the Aquascope and the microscopy data within the non-NA data decreases with increasing taxonomic resolution (**Tab.2**, with little difference between phylum and class level).

**Tab. 2:** Relative abundance (%) of different taxonomic levels from an integrated sample from Greifensee (Schröder 0-20m, 12.04.2023) measured with traditional microscopy and the Aquascope aggregated into different taxonomic levels. Linear model statistics where the Aquascope data (bioarea/L from ROI/sec) was fitted against the microscopy comparison (biovol/L from microscopy count). See also ***Tab. 1***.

| Taxonomic level | slope | p | Adjusted R2 | % NA in Aquascope data | % NA in microscopy data |
| --- | --- | --- | --- | --- | --- |
| phylum | 0.6473 | 0.012 | 0.3735 | 0 | 0 |
| class | 0.6507 | 0.0001 | 0.5257 | 0.2287 | 0 |
| order | 0.1942 | 0.0591 | 0.0704 | 0.6076 | 0.1254 |
| family | 0.188 | 0.0149 | 0.1179 | 0.7524 | 0.1725 |

As of July 2023, we can detect 29 phytoplankton taxonomic groups (i.e. general taxonomic categories, with 61 individual classifier categories) and 29 zooplankton taxonomic groups (i.e. general taxonomic categories, with 50 individual categories) across the two magnifications (**Fig. S1**, **Fig. S2**). With traditional microscopy, trained taxonomists can distinguish around 100 taxonomic groups, many at genus and even species level. Therefore, the Aquascope is unable to achieve the same taxonomic resolution that microscopic counts can provide. Other imaging techniques are known to have the same drawback, which is therefore not limited to the Aquascope. With the FlowCam for example, Alvarez et al. (2013) were able to distinguish 34 taxonomic groups, while a taxonomist was able to distinguish 144 taxonomic groups under the microscope. Nevertheless, most cantons report the abundance and biomass on phyla or class level to the general public (e.g. **BE**: (Guthruf, Maurer, and Zeh 2020); **AG**: (“Zustand Hallwilersee” n.d.); **ZH**: (Niederhauser et al. 2023)). Using imaging techniques, the continuation of those time series could be possible with a larger project to benchmark the different techniques.

Classifiers are only as good as their training data set and therefore, new taxa that enter a system, e.g. neobiota, might be missed because they are assigned to a pre-existing category. We therefore recommend adjusting and improving the ML classifiers regularly. Initially, trained taxonomists would need to build an expanded image library for new machines and study systems, but the training of a new ML classifier could build on the existing classifier for Greifensee. Once established, the minor improvements of the classifier would be significantly faster, especially because the existing ML classifier could be used to pre-sort the images for manual annotation.

### Sample processing speed

The Aquascope and the FlowCam have the potential to significantly speed up the enumeration of a plankton sample.

The number of species detected, and thus the accuracy in the representation of the sampled plankton community (including rare species), also relies a lot on the amount of sample that is used for enumeration and the number of particles counted (Rodríguez-Ramos et al. 2013). Usually, for microscopy counts, subsamples of 25, 50 or 100 mL, depending on the density of the sample, are sedimented into an Utermühl chamber. Then, a subset of all fields of view is counted in order to estimate the abundance of phytoplankton and microzooplankton. As a rule of thumb, 40 fields of view in two magnifications are enumerated. In the sample from Greifensee from 12.04.2023 that was used in this report, a total of 1554 particles were counted from a 25 mL sedimented subsample. A trained taxonomist needs approximately 1 to two days to count one sample.

In the Aquascope, within 5 minutes, a volume of 1.2 - 60mL (0p5x magnification) and 0.06 - 3 mL (5p0x magnification) is imaged with a frame rate of 1 per second (imaged volumes per frame from Merz et al. 2021). In order to enumerate the same volume as in traditional microscopy, one would have to image a phytoplankton sample for 41.67, 83.34 or 166 min under the assumption that the maximal theoretical imaged volume per frame is imaged in the 5p0x magnification. It took us 50 min to record 1270 ROI with the Aquascope on the Greifensee sample from 12.04.2023. As argued above, we would have to aim at a higher number of ROI in order to have a representative estimate of the plankton community. Nevertheless, it would take us no more than approximately two hours to process and analyse the sample that took about a day to process under the microscope. Data processing, once a pipeline is established, can then be done fully unsupervised and would likely not take more than a few minutes.

In comparison, the FlowCam usually runs at a speed between 0.03 mL/min (20x magnification) and 10 mL/min (2x magnification). Therefore, the time needed to enumerate a similar volume as in traditional microscopy, one would have to invest between 27 min (4x magnification, 25 mL total at 100 % efficiency, 0.9 mL/min) and 55 h (20x magnification, 100 mL total at 100 % efficiency, 0.03 mL/min). The advantage of the FlowCam is that the sample can be aspirated directly from a container through a tube and there is no additional supervision needed while the sample is running.

## Conclusion and Outlook

As shown in this report, there are large advantages in using automated imaging techniques for monitoring plankton. For the Aquascope, raw and processed data produced can be stored easily in a database, can be re-analysed over time as classification of plankton evolves, and can be shared. All software that is needed to deal with the data is open source and there is no profit-seeking company behind it. As a drawback, developments occur as a consequence to research projects, and there is no readily available solution at the moment to use the Aquascope as a lab instrument. On the other hand, the other image based plankton monitoring technique, the FlowCam, is an instrument designed for use in a laboratory. It has an annotation software implemented that is under licence. The extraction of the raw data from the software is not straightforward and there are no open source classifiers for the FlowCam images.

As of the moment, the Aquascope is not able to substitute long term monitoring data with the same taxonomic resolution and abundance estimation. There are a number of tests that would have to be done in a larger project in order to establish a working protocol from plankton monitoring with the Aquascope as a lab instrument. Namely:

– Establish a working lab adapter with a waterproof sealing that allows a constant flow rate (costs would lie around 2’500 CHF)
– Establish an improved colour transfer script, if necessary, to transfer images from any camera to match those used in the Greifensee classifier
– Either update or rebuild a classifier for multiple cameras and/or lakes
– Expand the calibration curves between ROI/sec and cells/mL for more species, including colonial phytoplankton, microzooplankton and multiple species of mesozooplankton. Additionally, assess whether this calibration is valid for different machines.
– Additional benchmarking of i) taxon-accumulation-curves, ii) relationships between microscopy and Aquascope data

To conclude, in comparison with traditional microscopy, automated imaging techniques can speed up the monitoring process significantly. In addition, the processing of a sample does not rely on the availability of a trained and experienced taxonomist which will be the limiting factor in the future. Overall, automated imaging techniques are powerful tools that pose a trade-off between i) traceability, individual-level traits, speed and independence from trained taxonomists and ii) taxonomic resolution and an expert’s insight. We show that with a larger benchmarking project (this applies for the Aquascope and the FlowCam), it might be possible to continue the long term time series on a coarse taxonomic level as usually reported by cantonal authorities to the general public. For this, and in addition with the development of a machine learning classifier, we recommend joining forces between cantons. In our opinion, the development of an automated imaging technique for plankton monitoring poses a large opportunity for collaboration on a federal level.

## Definitions and abbreviations

SPC: Scripps Plankton Camera
DSPC: Dual Scripps Plankton Camera (a dual-magnification dark field underwater microscope)
ML: machine learning
ROI: region of interest (single objects (i.e. a cell, a colony or a multicellular individual) generated by the underwater microscope)
Aquascope: synonym to DSPC
SPC-EAWAG: Aquascope installed in Greifensee. The ML classifier developed by Pomati, Baity Jesi and their teams is based on images from this instrument.
SPC-EAWAG2: secondary Aquascope in the lab.
Lake camera: synonym to SPC-EAWAG
Lab camera: synonym to SPC-EAWAG2
0p5x: 0.5x magnification targeting mesozooplankton, microzooplankton and large phytoplankton colonies
5p0x: 5x magnification targeting phytoplankton and microzooplankton
Utermöhl chamber: standard (phyto-)plankton sedimentation and counting chamber usually used with inverted microscopy (Utermöhl 1931)

## Acknowledgement

Marco Baity Jesi and Sreenath Kyathanahally for the colour transformation script that was developed especially for this project.

Lukas De Ventura, Kristina Rehberger, Katrin Gutruf, Vincent Maurer and Silvana Käser for samples, data, discussions and support.

We thank Michael Koss (WVZ) for showing us the FlowCam and sharing example images from their instrument (see Fig. S3).

We thank the cantons of Aargau, Bern, Luzern, Fribourg, Neuchâtel, Vaud and Zurich for financing this project.

## Supplementary

### Images from the Aquascope

**Fig. S1:**
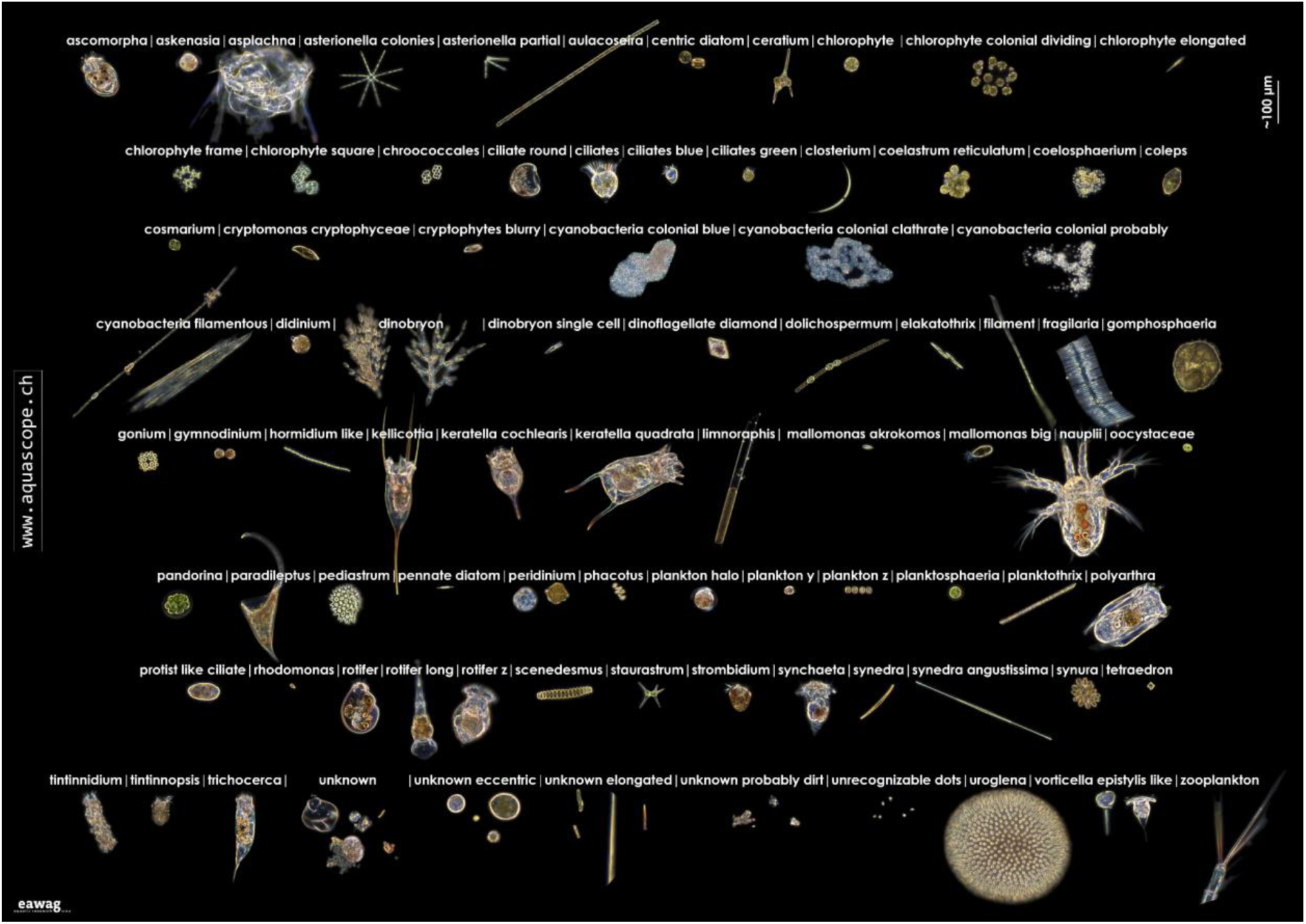
All categories (taxa) we can distinguish and automatically classify in the 5p0x magnification of the Aquascope. Categories are built for the use of the ML classifier and based on optical dissimilarity which enhances classification. Some categories are later summed into general taxonomic categories based on taxonomy.

**Fig. S2:**
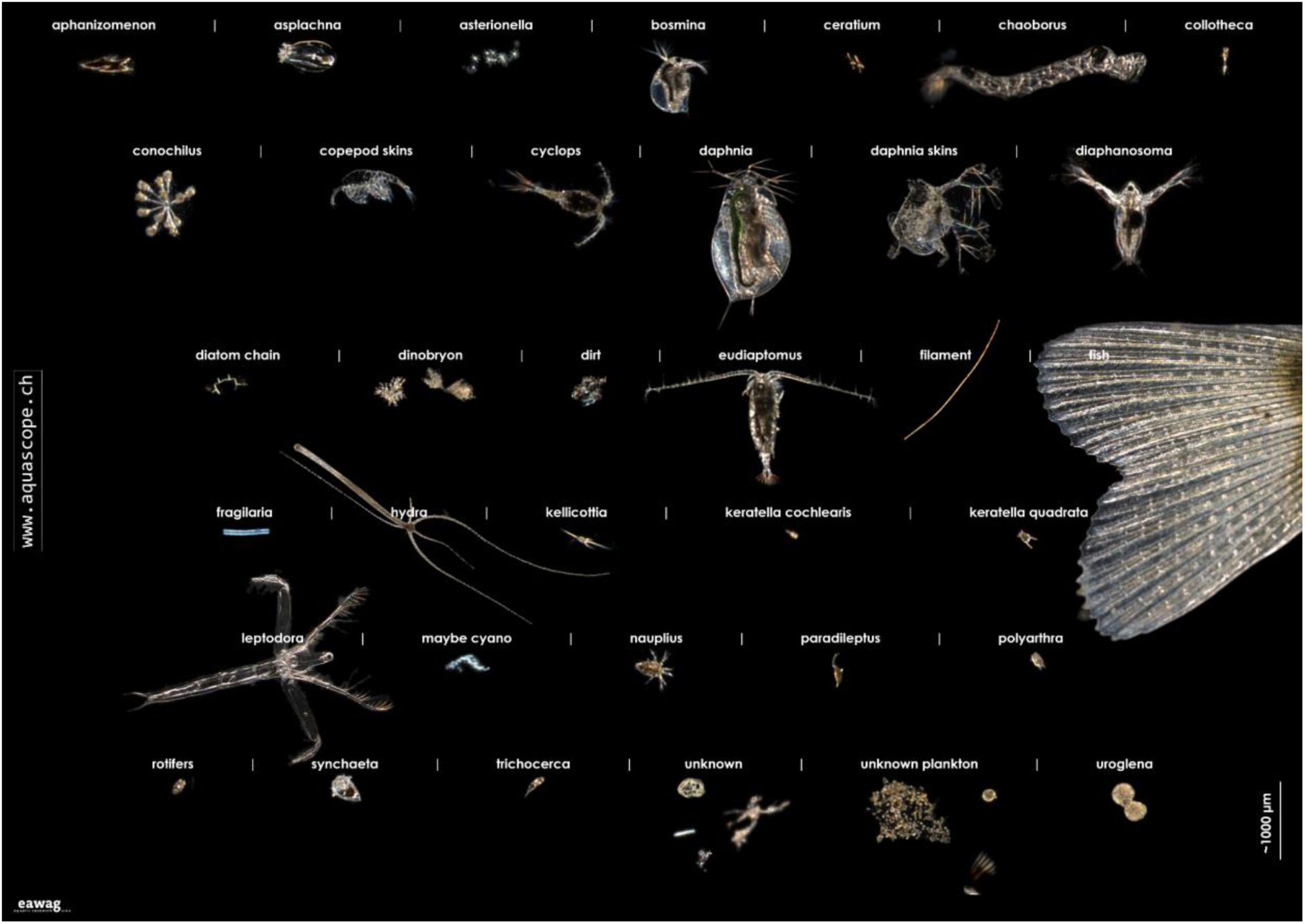
All categories (taxa) we can distinguish and automatically classify in the 0p5x magnification of the Aquascope. Categories are built for the use of the ML classifier and based on optical dissimilarity which enhances classification. Some categories are later summed into general taxonomic categories based on taxonomy.

**Fig. S3:**
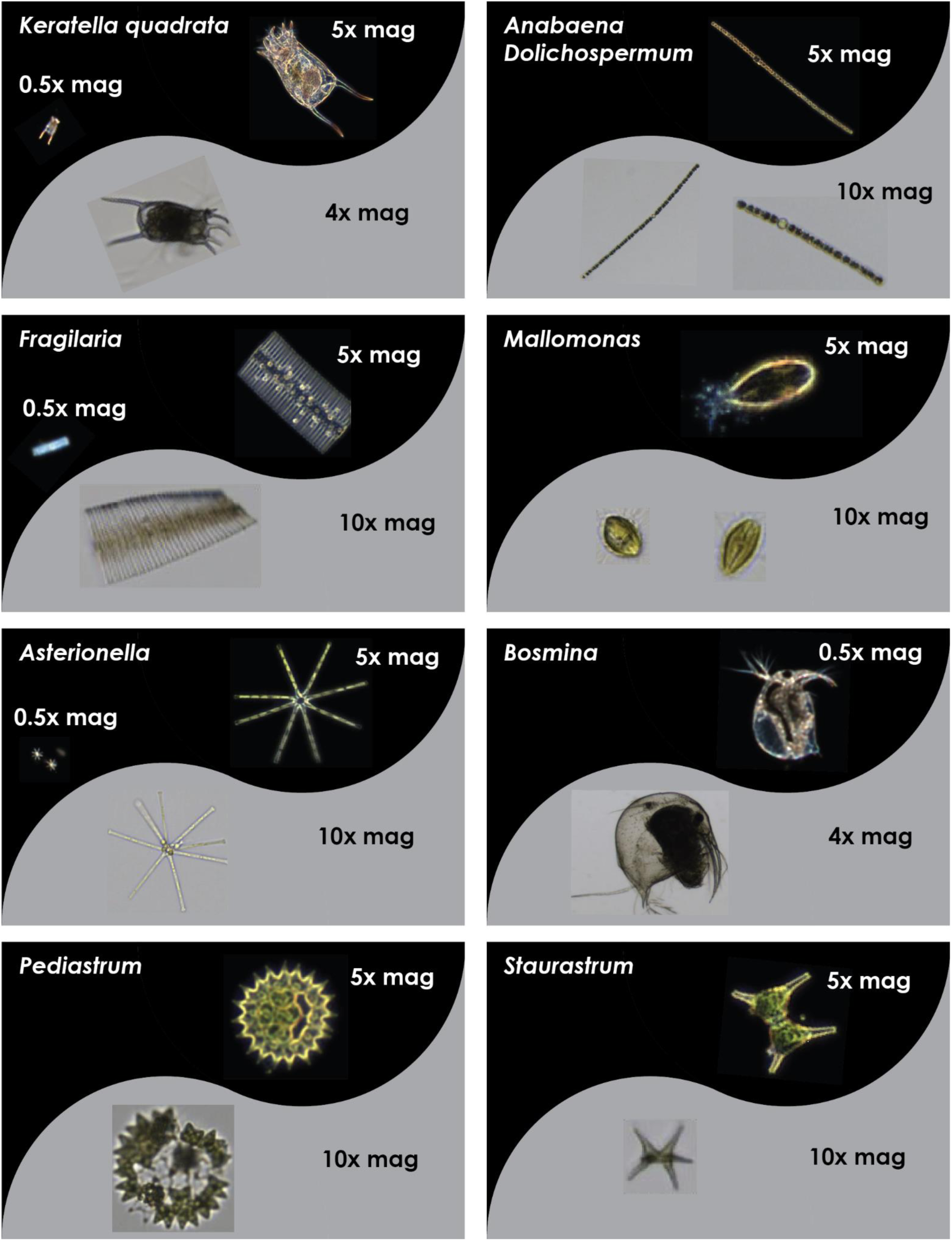
Comparison of the images generated by the Aquascope and by the FlowCam (images by WVZ, Michael Koss). Images are shown with the magnification in which they are taken. On black background, images from the Aquascope. On grey background, images from the FlowCam.

### Supplementary plots

#### Colour legends

**Fig. S4:**
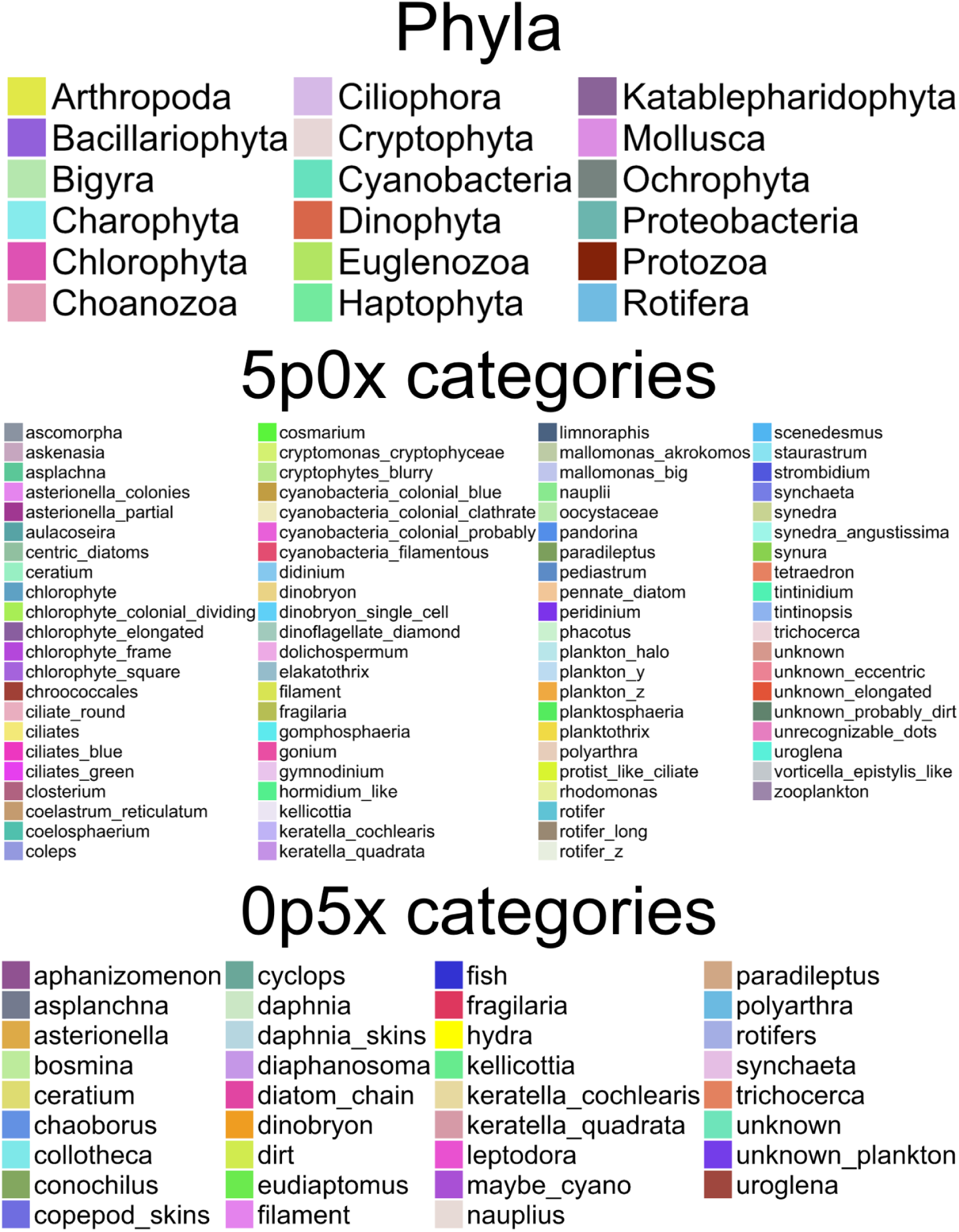
Colour legend corresponding to all plots in the report. The colours are the same in all plots with taxonomic information. The two magnifications from the Aquascope have separated colour legends.

#### Data from Greifensee

**Fig. S5:**
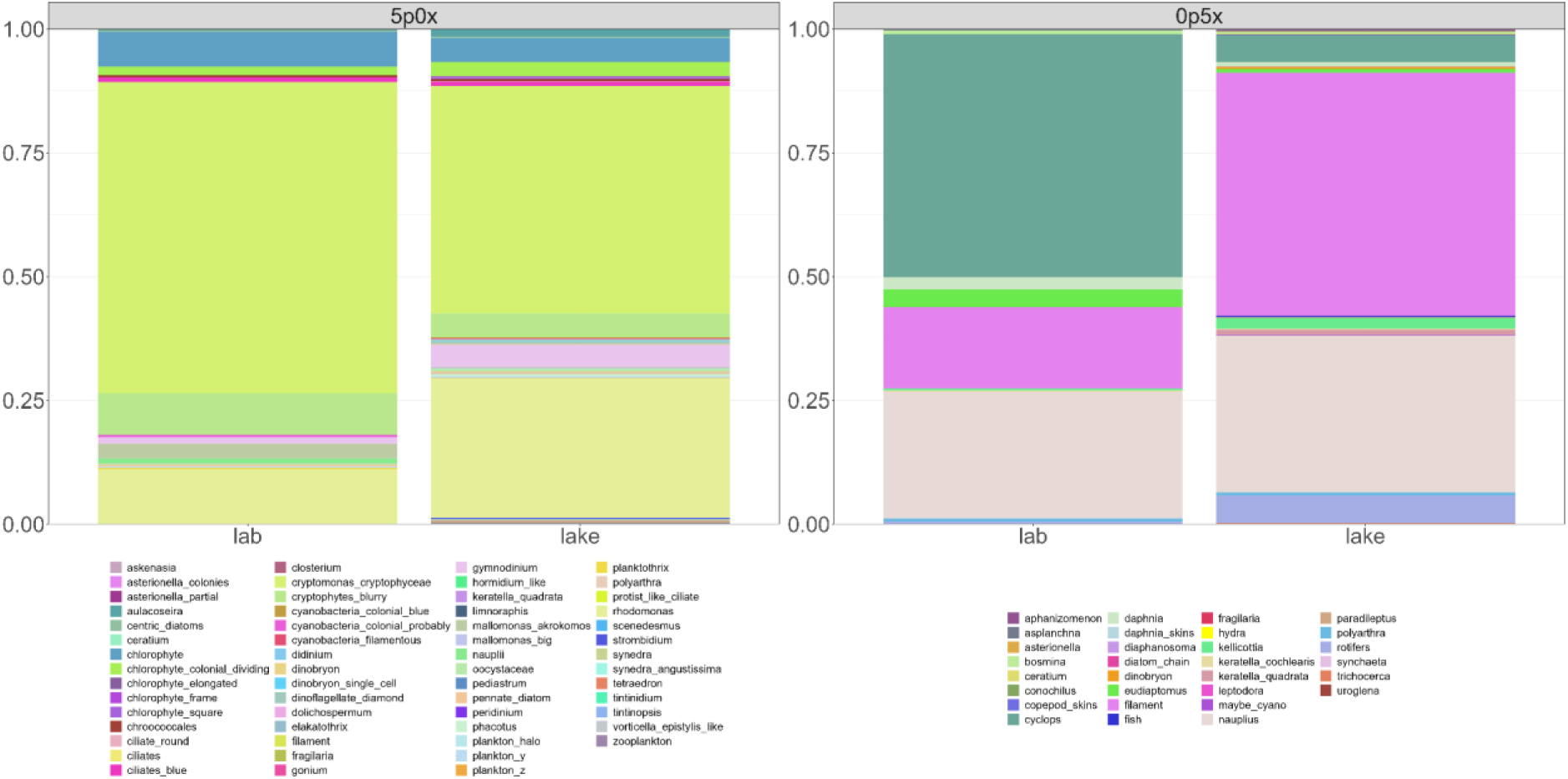
Comparison between a 3m sample from Greifensee (Niskin, 23.03.2023, “lab” column) and the images taken at 3m with the lake camera (“lake” column) on the same day (23.03..2023 00:00 - 23:59). Separated by magnification into two side-by-side plots (left: 5p0x magnification, right: 0p5x magnification). Displayed are the relative abundances of the mean ROI/sec across all lab camera runs and the ROI/sec from the lake camera. Unknown classes (unknown, unknown_eccentric, unknown_probably_dirt, unrecognizable_dots, unknown_plankton, bubbles) in the 0p5x magnification were removed.

**Fig. S6:**
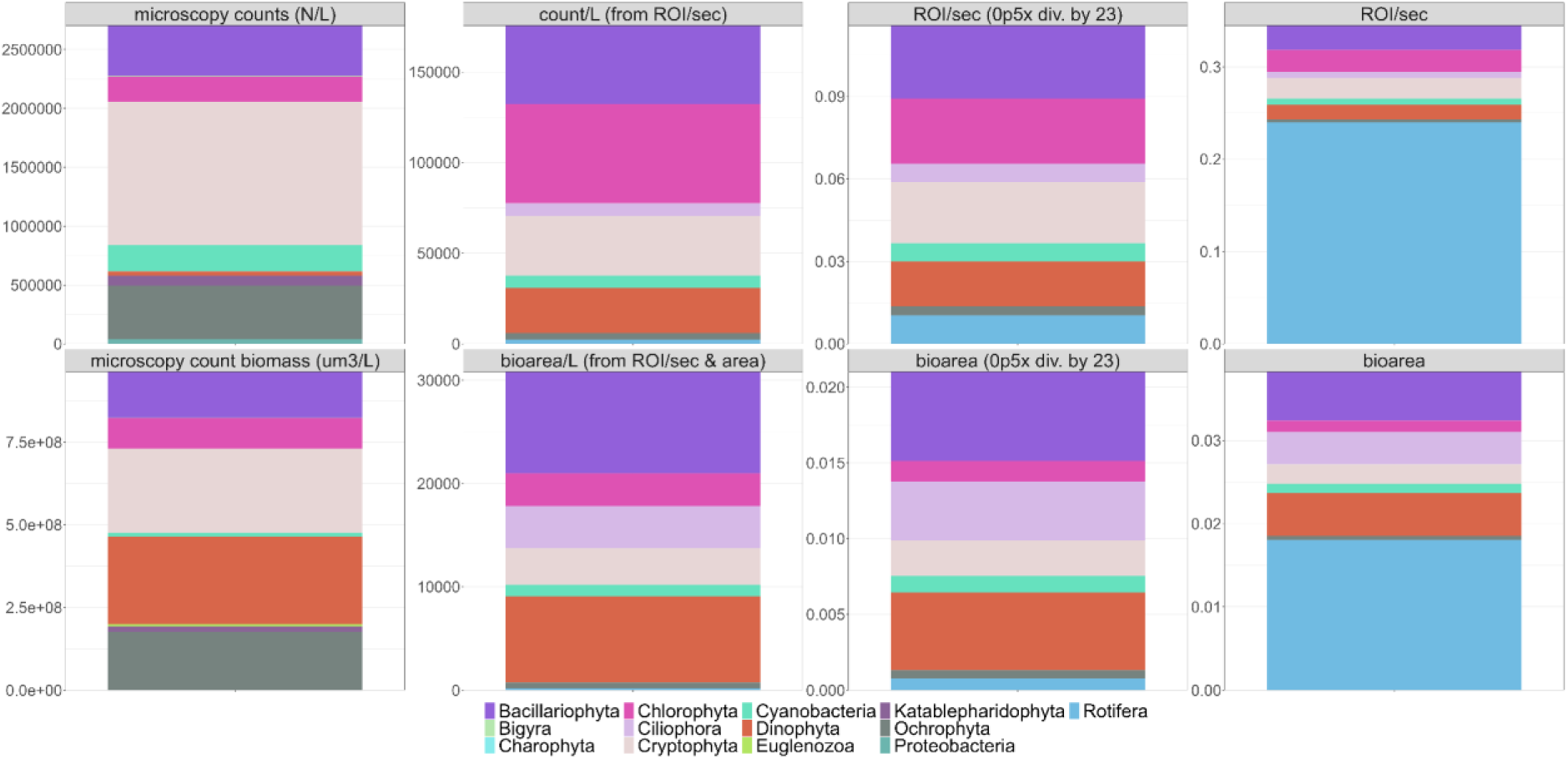
Integrated sample from Greifensee (Schröder 0-20m, 12.04.2023) measured with traditional microscopy and the Aquascope aggregated into phyla. **Top row**: abundance (from left to right: microscopy cells/L, cells/L calculated from ROI/sec, raw ROI/sec with abundances from the 0p5x magnification divided by 23, raw ROI/sec). **Bottom row**: biomass (from left to right: biomass/L (μm^3^/L) from the microscopy counts, bioarea/L (mm^2^/L) calculated from the total bioarea and the cells/L, raw bioarea (mm^2^) with bioarea from the 0p5x magnification divided by 23, raw bioarea (mm^2^)).

#### Data from Canton Bern

**Fig. S7:**
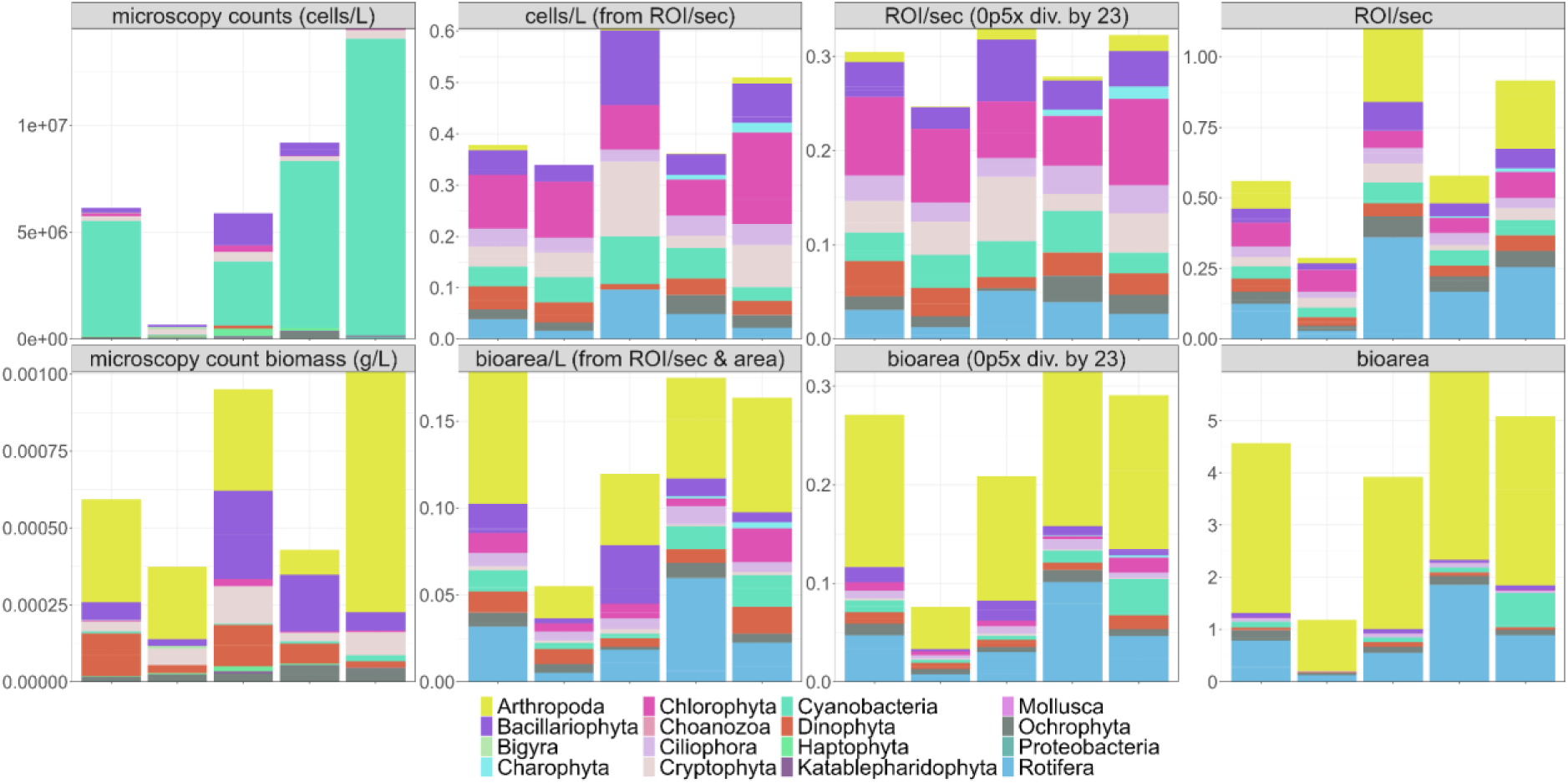
Samples from Canton Bern measured with traditional microscopy and the Aquascope aggregated into phyla. The samples come from different lakes, at different times of the year: Lake Thun (Aug, Nov), lake Brienz (Sep), Biel (Oct), and Murten (Dec). **Top row**: abundance (from left to right: microscopy cells/L, cells/L calculated from ROI/sec, raw ROI/sec with abundances from the 0p5x magnification divided by 23, raw ROI/sec). **Bottom row**: biomass (from left to right: biomass/L (g/L) from the microscopy counts, bioarea/L (mm^2^/L) calculated from the total bioarea and the cells/L, raw bioarea (mm^2^) with bioarea from the 0p5x magnification divided by 23, raw bioarea (mm2)).

**Fig. S8:**
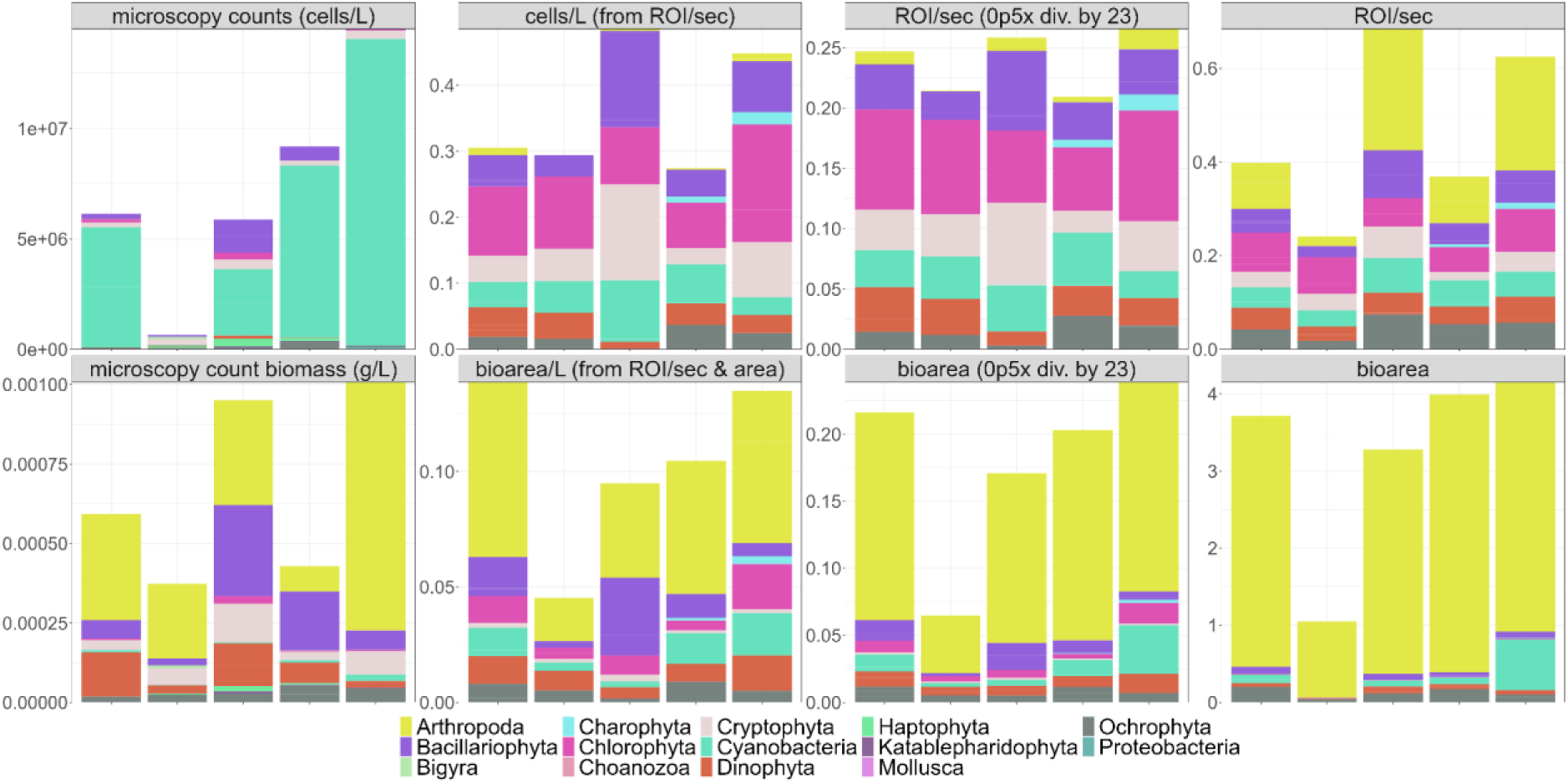
Samples from Canton Bern measured with traditional microscopy and the Aquascope aggregated into phyla. The samples come from different lakes, at different times of the year: Lake Thun (Aug, Nov), lake Brienz (Sep), Biel (Oct), and Murten (Dec). Here, Rotifera and Ciliophora have been removed as they are not counted by the canton of Bern. **Top row**: abundance (from left to right: microscopy cells/L, cells/L calculated from ROI/sec, raw ROI/sec with abundances from the 0p5x magnification divided by 23, raw ROI/sec). **Bottom row**: biomass (from left to right: biomass/L (g/L) from the microscopy counts, bioarea/L (mm^2^/L) calculated from the total bioarea and the cells/L, raw bioarea (mm^2^) with bioarea from the 0p5x magnification divided by 23, raw bioarea (mm^2^))

#### Data from Hallwilersee

**Fig. S9:**
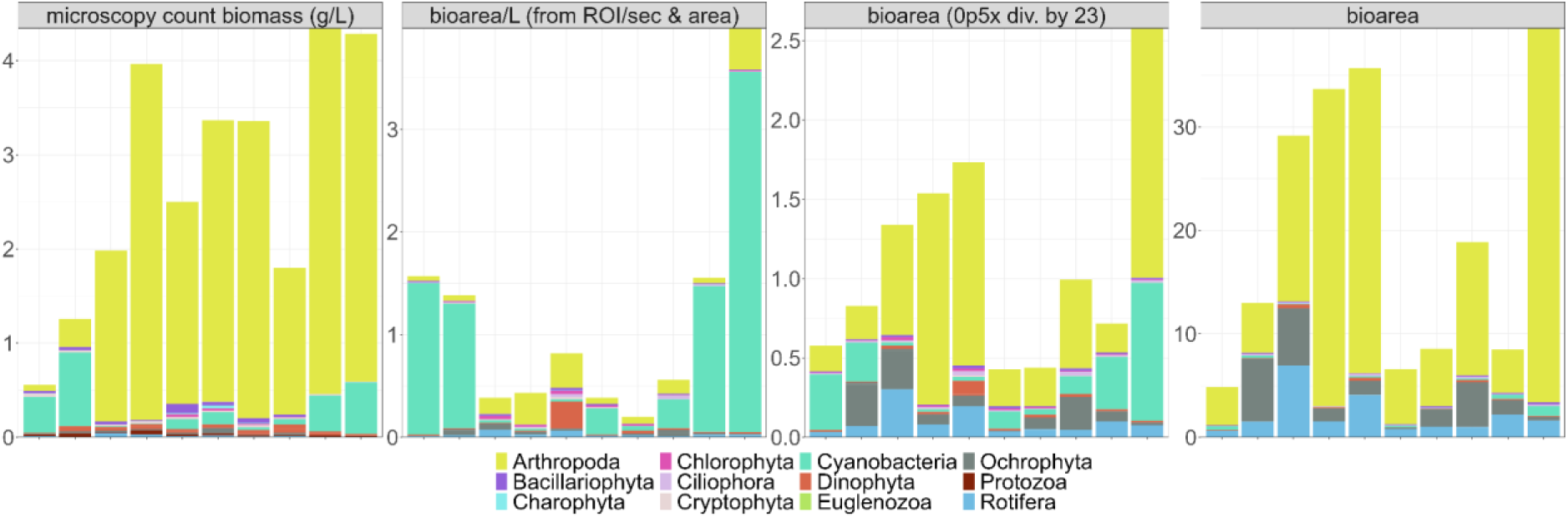
Samples from Hallwilersee measured with traditional microscopy and the Aquascope aggregated into phyla. The samples were taken every month in 2022 starting in March. Bars are arranged from March to December from left to right in each plot. Displayed is the biomass (from left to right: biomass/L (g/L) from the microscopy counts, bioarea/L (mm^2^/L) calculated from the total bioarea and the cells/L, raw bioarea (mm^2^) with bioarea from the 0p5x magnification divided by 23, raw bioarea (mm^2^)). Note that Arthropoda were sampled differently for microscopy (twin-net to 43m instead of 0-13m integrated sample that was used for the rest of the microscopy counts and the Aquascope measurements.

**Fig. S10:**
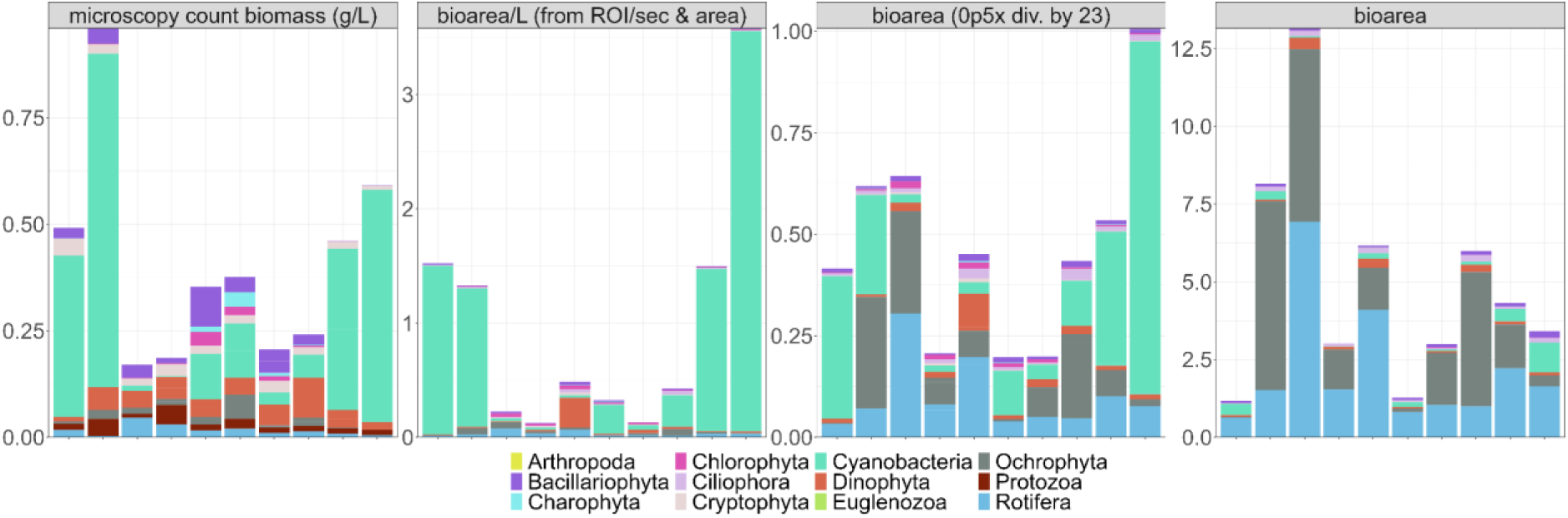
Samples from Hallwilersee measured with traditional microscopy and the Aquascope aggregated into phyla. The samples were taken every month in 2022 starting in March. Bars are arranged from March to December from left to right in each plot. Displayed is the biomass (from left to right: biomass/L (g/L) from the microscopy counts, bioarea/L (mm^2^/L) calculated from the total bioarea and the cells/L, raw bioarea (mm^2^) with bioarea from the 0p5x magnification divided by 23, raw bioarea (mm^2^)). Arthropoda were removed because they were sampled differently for microscopy (twin-net to 43m instead of 0-13m integrated sample that was used for the rest of the microscopy counts and the Aquascope measurements.

**Fig. S11:**
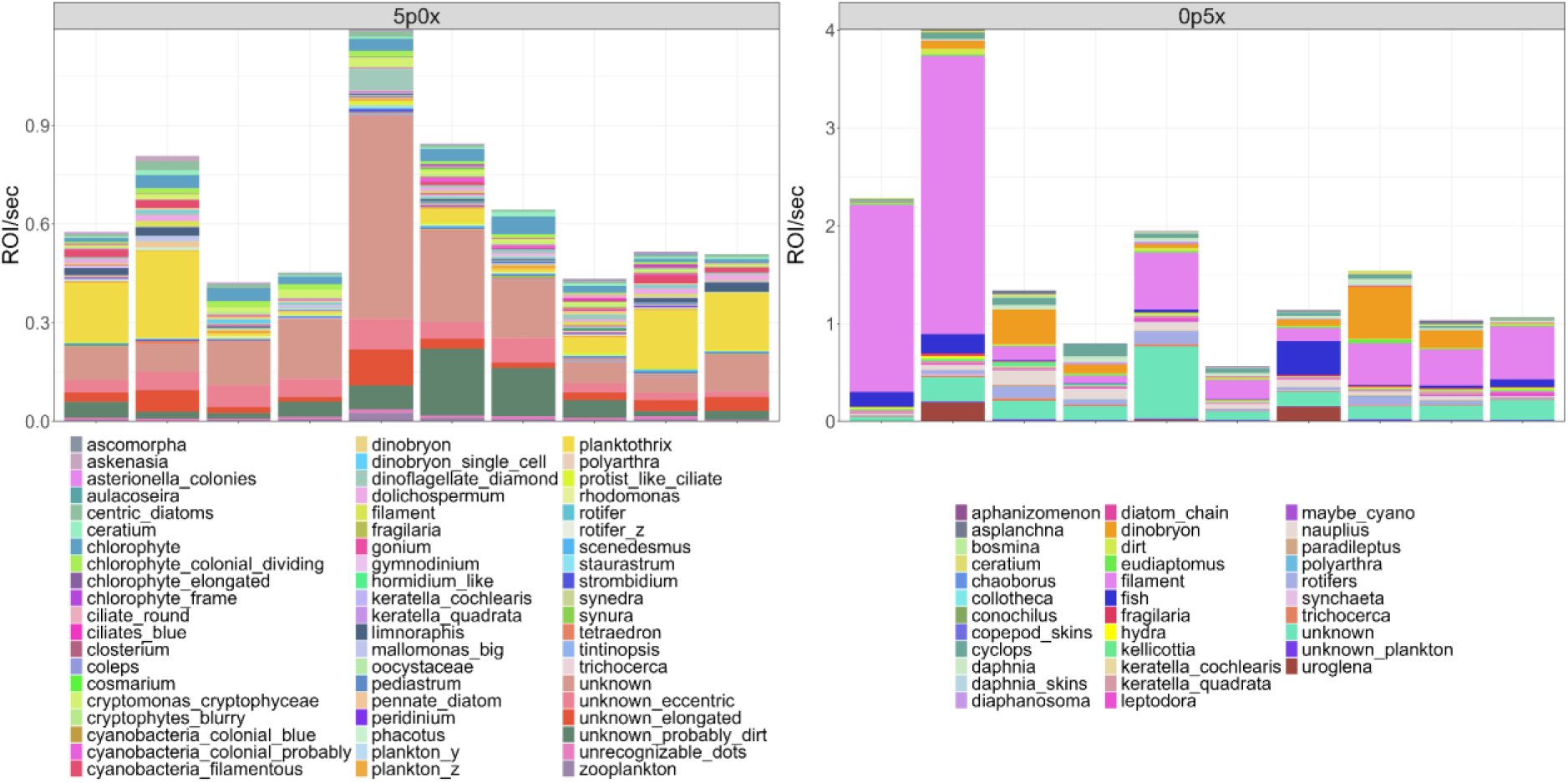
Samples from Hallwilersee measured with the Aquascope. The samples were taken every month in 2022 starting in March. Bars are arranged from March to December from left to right in each plot. Separated by magnification into two side-by-side plots (left: 5p0x magnification, right: 0p5x magnification). Displayed are the abundances (mean ROI/sec across all lab camera runs) of each Aquascope category.

### Supplementary tables

**Tab. S1:** One-sample T-test on the ROI/sec we expect from a perfect dilution series based on the mean ROI/sec across eight replicates in each magnification at 100% dilution. H0: mean = theoretical mean (μ). At a p < 0.05, the observed mean is significantly different to the theoretical mean. This is only the case for the 0p5x magnification at 1:7 dilution (12.5%).

| magnification | dilution | p | Theoretical mean ($\mu$) | sample size |
| --- | --- | --- | --- | --- |
| 0p5x | 50 % (1:1) | 0.4003 | 0.229479 | 4 |
|  | 25 % (1:3) | 0.6861 | 0.11474 | 3 |
|  | 12.5 % (1:7) | 0.0183 | 0.05737 | 3 |
| 5p0x | 50 % (1:1) | 0.7278 | 0.102188 | 4 |
|  | 25 % (1:3) | 0.3284 | 0.051094 | 3 |
|  | 12.5 % (1:7) | 0.2679 | 0.025547 | 3 |

**Tab. S2:**
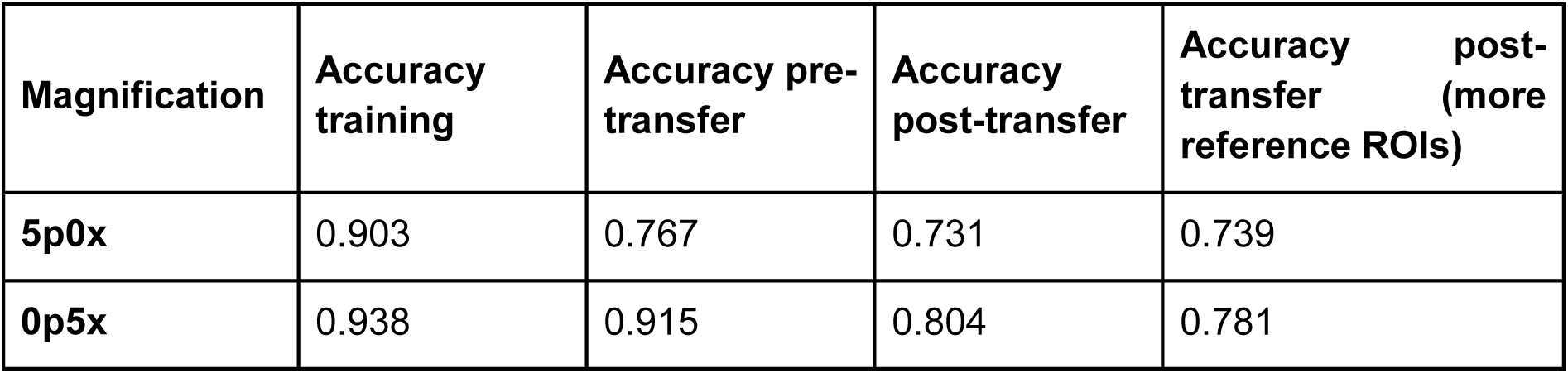
Accuracy of the classifier from training (https://github.com/kspruthviraj/Plankiformer) and on a sample from Greifensee (12.04.2023), imaged in the lab. The accuracy of the 0p5x and the 5p0x magnification was assessed before and after using the colour transfer script. For the colour transfer, we used 1727 ROIs or 5178 ROIs from the 0p5x magnification and 6219 ROIs or 13287 ROIs from the 5p0x magnification as a reference.

**Tab. S3:**
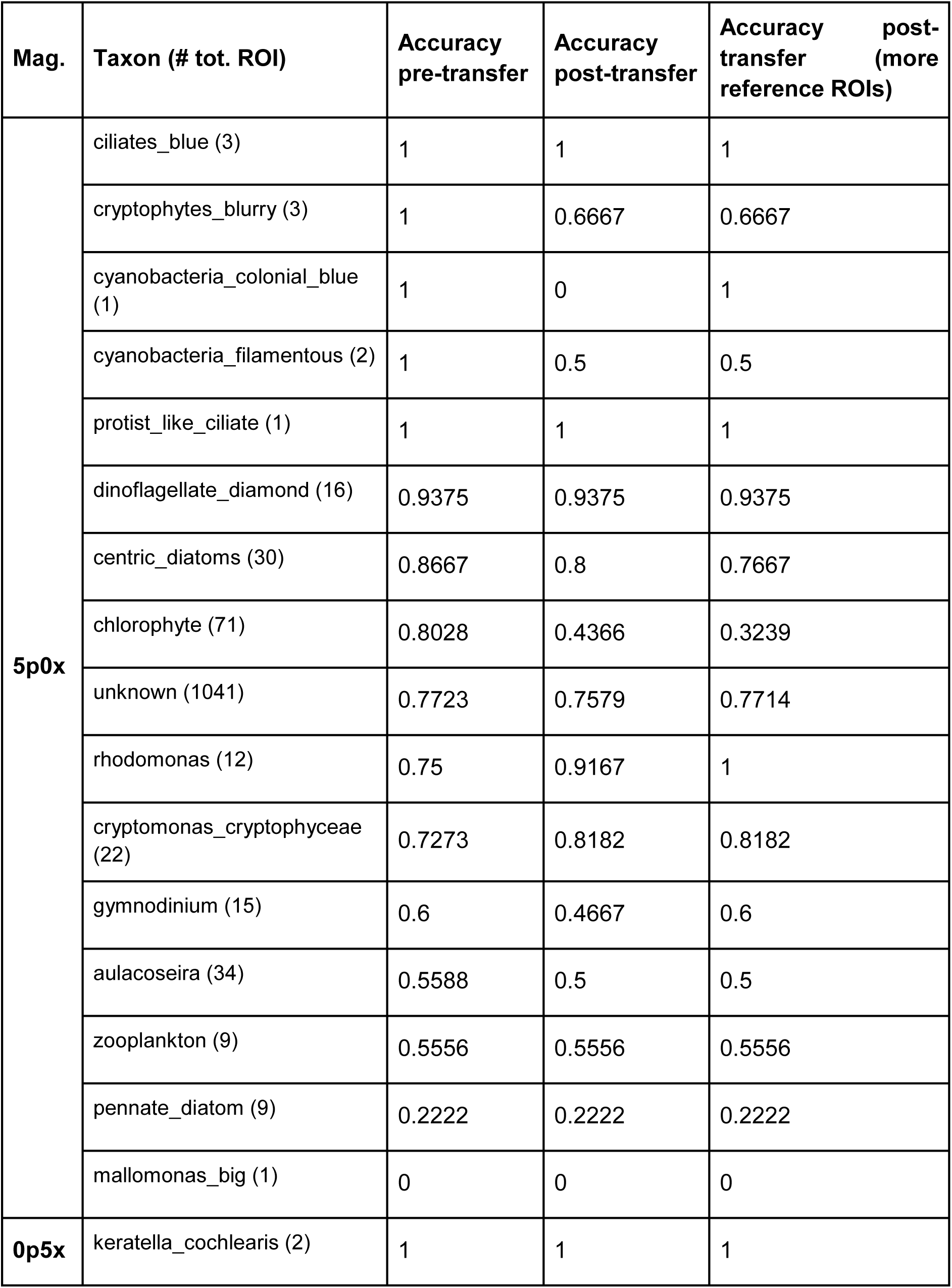
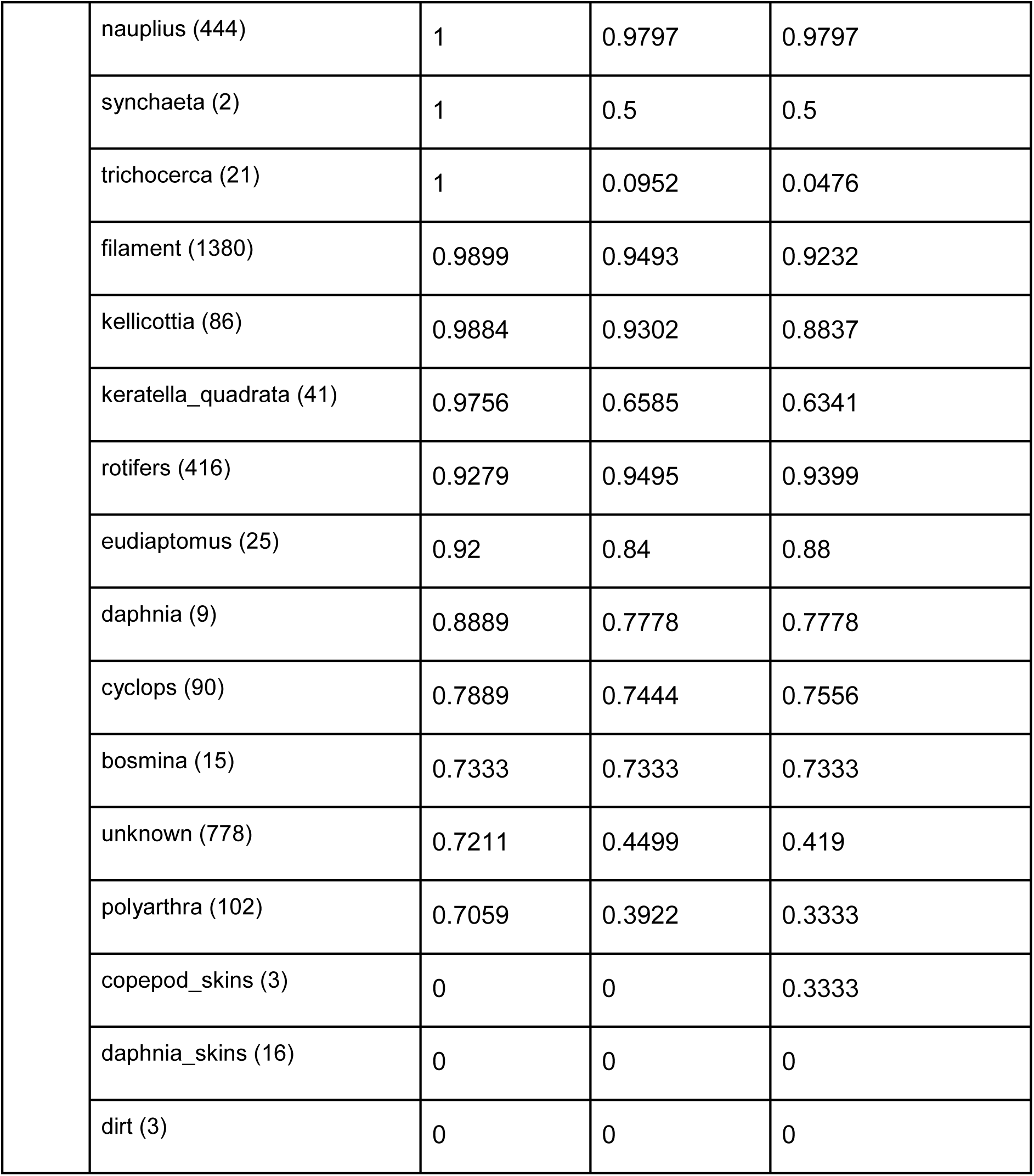
Accuracy of the classifier on a sample from Greifensee (12.04.2023), imaged in the lab. The accuracy of each taxon of the 0p5x and the 5p0x magnification was assessed before and after using the colour transfer script. For the colour transfer, we used 1727 ROIs or 5178 ROIs from the 0p5x magnification and 6219 ROIs or 13287 ROIs from the 5p0x magnification as a reference.

